# A first-in-kind MAPK13 inhibitor corrects epithelial stem cell reprogramming and muco-obstructive lung disease

**DOI:** 10.1101/2024.08.21.608990

**Authors:** Yong Zhang, Kangyun Wu, Dailing Mao, Courtney A. Iberg, Huiqing Yin-Declue, Kelly Sun, Hallie A. Wikfors, Shamus P. Keeler, Ming Li, Deanna Young, Jennifer Yantis, Erika C. Crouch, Joshua R. Chartock, Zhenfu Han, Kay O. Broschat, Derek E Byers, Steven L. Brody, Arthur G. Romero, Michael J. Holtzman

## Abstract

The stress kinase MAPK13 (aka p38δ-MAPK) is an attractive entry point for therapeutic intervention because it regulates the structural remodeling that can develop after epithelial injury in the lung and likely other tissue sites. However, a selective, safe, and effective MAPK13 inhibitor is not yet available for experimental or clinical application. Here we identify a first-in-kind MAPK13 inhibitor using structure-based drug design combined with a screening funnel for cell safety and molecular specificity. In a mouse model of severe respiratory viral infection, treatment with this inhibitor (formulated as NuP-4A for intravenous use or Nu4-B for inhaled delivery) did not influence recovery from acute infectious illness, but still down-regulated basal-epithelial stem cell (basal-ESC) hyperplasia/metaplasia and in turn airway inflammation, mucus production, and pathophysiology biomarkers of chronic lung disease. Treatment prevented and reversed disease readouts equivalently to *Mapk13* gene-knockout, and this benefit persisted after stopping treatment as a sign of disease modification. Further, NuP-4 treatment even at pM levels directly blocked basal-ESC reprogramming endpoints in organoid and cell-culture models derived from non-disease control and asthma subjects. The results thereby provide a new tool compound and drug candidate for basal-ESC reprogramming towards muco-obstructive lung diseases like asthma and any overlap with COPD and related diseases that depend on overactivity of MAPK13.

**New and noteworthy:** This study identifies to our knowledge a first highly selective and potent small-molecule inhibitor (designated NuP-4) to target stress kinase MAPK13. The results also demonstrate that NuP-4 treatment preserves homeostatic defense and repair but prevents excessive basal-ESC responses to respiratory viral infection and thereby attenuates key biomarkers of chronic lung disease in mouse and human models. The findings provide guidance for precise therapeutic application in asthma and related diseases that are triggered by respiratory viruses and other airborne hazards. The tissue distribution of MAPK13 implies related applications at other epithelial sites.

## Introduction

The family of mitogen-activated protein kinases (MAPKs), and in particular the subgroup of p38-kinases (MAPK11-14 aka p38α-δ-MAPK), are implicated in a broad set of experimental models of disease and the corresponding human diseases (1, 2). These studies led to a corresponding array of drug development programs directed at MAPK targets, particularly MAPK14 (aka p38α-MAPK) as the first and best studied member of the MAPK family (3–11). Despite this effort, no inhibitors for MAPK14 have been approved for clinical application. The basis for failure in clinical trials range from suboptimal efficacy and safety to difficulties in target validation, biomarker application, and disease modification. Nonetheless, MAPK14 and related kinases continue to be an attractive target for ongoing drug discovery and development programs.

In that context, the previously undrugged MAPK13 (aka p38δ-MAPK) is implicated in diseases that include diabetes, sepsis, neurodegeneration, and carcinoma (12–17). Given this diversity, we questioned whether MAPK13 might represent a more fundamental control over cellular programming and reprogramming. Indeed, our earlier and ongoing studies identified MAPK13 as a therapeutic target in chronic lung disease. This idea derived from a MAPK13 requirement for mucus production in human basal-epithelial stem cell (basal-ESC) models and a corresponding upregulation of MAPK13 expression and activation in lung tissue samples from COPD patients (18). In follow up, recent work demonstrates a MAPK13 requirement in vivo. Thus, the long-term basal-ESC reprogramming and muco-obstructive lung disease that develops after respiratory viral infection is markedly and safely attenuated in *Mapk13* gene-knockout mice (19). In addition, *MAPK13* gene-deficiency significantly down-regulated basal-ESC hyperplasia in primary mouse and human cell models as evidence of a direct role for MAPK13 in epithelial cell function (19). These findings provided a pathway to practical application since epithelial stem cell-orchestrated inflammation and mucus production are linked to morbidity and mortality in chronic muco-obstructive lung diseases such as asthma and COPD (18, 20–22).

The present study therefore addresses the goal of discovering and developing a highly selective MAPK13 inhibitor to solve the problem of epithelial stem cell reprogramming towards chronic disease. Previous studies by our group could not specifically address this issue given the broader specificity of first- and second-generation inhibitors and the absence of studies in a model of basal-ESC reprogramming that might exhibit long-term inflammation and tissue remodeling (18, 23). In contrast, the present study arrives at a third-generation MAPK13 inhibitor that is remarkably selective compared to previous compounds from our lab and others (11, 18, 23, 24). In addition, the proof-of-concept is developed in a suitable model for basal-ESC reprogramming for persistent hyperplasia and mucous metaplasia after viral infection and likely other triggers for disease at epithelial barrier sites. Together, the present experimental approaches provide a previously unavailable tool for MAPK13 blockade and proof-of-concept for titrating down-regulation of this kinase target as a disease-modifying strategy.

## Results

### Structure-based drug design yields a selective MAPK13 inhibitor NuP-4

Given the benefits of MAPK13 deficiency in our models of lung disease, we addressed the need for a small-molecule inhibitor that was selective for MAPK13 versus the historical focus on MAPK14 (24) or in our case MAPK13-14 (24). Accordingly, we extended a drug-design strategy to fully modify a selective MAPK14-inhibitor parent compound (BIRB-796, NuP-43 in our chemical library (as depicted in **Figure 1A**) to better fit into the left-hand hinge region, ATP-binding pocket, and right-hand allosteric pocket of MAPK13 as defined in our high-resolution X-ray crystal structure (18, 23, 25, 26). In previous work, we produced first-generation compounds with a modest (3-7-fold) increase in MAPK13 inhibitory activity (18) and then second-generation compounds, including NuP-3 that improved MAPK13 inhibition (130-fold) compared to the parent compound NuP-43 (23). However, these compounds also maintained similar MAPK14 inhibition. Given our new evidence for MAPK13 function alone in vivo and ex vivo, we used our expanded library of chemical analogs (n=520) as recently published (27) to discover a more selective MAPK13 inhibitor. These analogs entered a screening funnel that included a primary screen for suitable chemical properties (molecular weight, Lipinski and Veber physical chemistry structure criteria, partition coefficient, and topological polar surface area) followed by a secondary screen for single-dose inhibition of MAPK13 and MAPK14 enzyme activities, a tertiary screen for cell toxicity based on transepithelial electrical resistance (TEER) as a readout for monolayer integrity in primary-culture human tracheobronchial epithelial cell (hTEC) cultures, and a full dose-response for MAPK13 enzyme inhibition (**Figure 1B**).

**Fig. 1.**
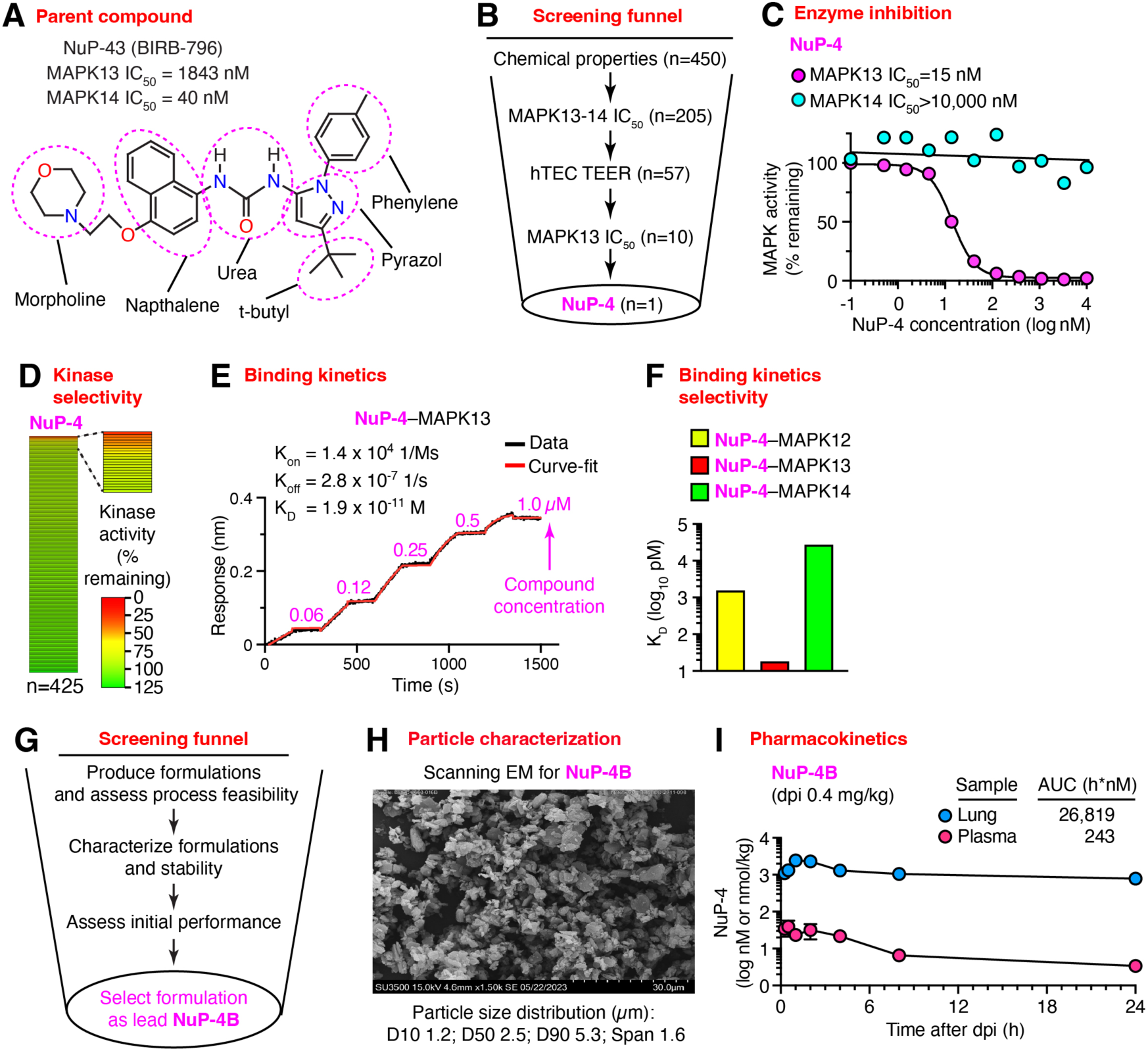
Structure-based drug design yields a lead drug candidate MAPK13 inhibitor (NuP-4). **A,** Chemical structure for parent compound NuP-43 (BIRB-796) as a starting point for modifications to increase inhibition of MAPK13. **B,** Screening funnel steps for chemical compound library using MAPK13-14 enzyme assays, trans-epithelial electrical resistance (TEER) for human tracheobronchial epithelial cell (hTEC) monolayer integrity for cell toxicity, and additional MAPK13 assays. **C,** Enzyme inhibition activities for NuP-4 based on MAPK13 and MAPK14 phosphorylation of kinase substrate. **D,** Heat map for NuP-4 selectivity against a 425-kinase panel. **E**, Biolayer interferometry (BLI) analysis for NuP-4–MAPK13 binding. **F**, Corresponding BLI-derived K_D_ for NuP-4 binding to MAPK12,13,14. **G**, Screening funnel steps for development of a formulation suitable for dpi with selection of NuP-4B. **H**, Particle characterization using scanning electron microscopy (EM) and resulting particle size distribution after jet-milling. **I**, Pharmacokinetic analysis of lung and plasma levels of NuP-4B (assayed as NuP-4) using dpi delivery to mice.

This process arrived at a lead drug candidate designated NuP-4 with a 122-fold increase in MAPK13 inhibition and >250-fold decrease in MAPK14 inhibition compared to the parent compound NuP-43 (**Figure 1C**). NuP-4 also exhibited a DFG-out binding mechanism that provides favorable slow-dissociation kinetics and consequent high potency and long duration of action of a Type II kinase inhibitor (18, 28, 29). This mechanism fits with structural modeling (30, 31). Further, NuP-4 demonstrated markedly enhanced selectivity for MAPK13 in a 425-kinase inhibitor screen compared to parent compound NuP-43 or MAPK13-14 inhibitor NuP-3 (**Figure 1D**). Notably, we also found 200-fold higher binding (lower K_D_) for NuP-4 to MAPK13 compared to even its closest homologue MAPK12 based on a remarkably long K_off_ rate (**Figure 1E,F**). Together, this data supports high potency and specific on-target activity of NuP-4, which fits with our analysis of MAPK13 expression and function as noted below in preclinical proof-of-concept (POC) and safety pharmacology and toxicology. Together, the data support strong and specific on-target activity for NuP-4. These characteristics translated to rat toxicology studies wherein dose-range finding to 60 mg/kg iv for 14 d caused no adverse effects or significant abnormalities in clinical laboratory values and markedly higher levels in lung tissue than plasma (**Supplemental Figure 1A-F**).

Initial work optimized NuP-4 solubility for intravenous (iv) delivery and arrived at the HBr salt (NuP-4A) in 2-hydroxypropyl-β-cyclodextrin excipient. Given the proposed application to lung disease, we launched a separate campaign for inhaled delivery. Indeed, NuP-4 was selected to meet all of the recommendations for inhaled dosing (32) by: (i) lowering intrinsic aqueous solubility (0.85 µg/ml); (ii) limiting dissolution rate (exhibiting slow kinetic solubility); (iii) lowering permeability for intestinal absorption (P_app_=1×10^-6^ cm/sec with efflux ratio=2.3); (iv) increasing lung tissue affinity by modulating compound lipophilicity (logP=5.1); (v) optimizing compound kinetic off-rate from the target (K_off_=multi-hour); and (vi) mechanism-specific approaches that are also target-specific as noted above in lead discovery. Subsequent work completed micronization feasibility and solid form selection screening (**Figure 1G**). A salt of NuP-4 (designated NuP-4B) was identified as the lead with suitable particle characteristics for jet-mill micronization (**Figure 1H**). Further, NuP-4B dry powder inhalation (dpi) in mice provided 110-fold higher levels in lung compared to plasma and duration consistent with once per day (qd) therapeutic dosing (**Figure 1I**). In addition, NuP-4B dpi showed similar pharmacokinetics and no observed adverse effects up to a single maximal feasible dose (MFD) of 7 mg/kg or multiple daily dosing up to 2 mg/kg for 7 d in rat toxicology studies (**Supplemental Figure 1G**).

### MAPK13 inhibitor NuP-4A prevents post-viral lung disease

We next determined whether NuP-4 might provide MAPK13 blockade and therapeutic benefit in vivo, taking advantage of our mouse model of lung disease that develops and persists after respiratory viral infection. For this model, NuP-4 was formulated as the corresponding salt (designated NuP-4A) in excipient cyclodextrin to achieve compound solubility suitable for parenteral dosing. Using this formulation, pharmacokinetic (PK) analysis showed that NuP-4A levels (assayed as free base NuP-4) were consistently increased in target lung tissue compared to plasma when given by iv or intraperitoneal (ip) injection to mice (**Figure 2A**) and rats (**Supplemental Figure 1F**). In the mouse model, NuP-4A dosing achieves 9-32-fold higher area-under-the-curve (AUC) in lung tissue compared to plasma consistent with high uptake and target affinity in lung. These lung tissue levels were maintained for at least 8 h at a concentration (>100 nM) that was effective for fully blocking MAPK13 enzyme activity (**Figure 2A** and **Figure 1C**). In that context, we proceeded with a dose of 4 mg/kg given ip twice per day (bid), and to approximate clinical treatment conditions, we initiated treatment at 5 d after infection and assessed animals at 49 d when disease phenotype is maximal (33–35). Under this protocol (as diagrammed in **Figure 2B**), we showed that the usual acute illness and associated weight loss were slightly improved in mice treated with NuP-4A (**Figure 2C**). Further, the usual progression to muco-obstructive lung disease was markedly attenuated in NuP-4A-treated mice. Thus, the typical increase in MAPK13^+^KRT5^+^ basal-epithelial cells in remodeling regions was significantly diminished with NuP-4A-treatment compared to vehicle control at 49 d after SeV infection (**Figure 2D,E**). In concert with control of basal cell overgrowth, levels of PAS-hematoxylin staining were also attenuated in NuP-4A-treated mice (**Figure 2F**). Treatment levels of MAPK13, KRT5, PAS, and hematoxylin staining were each confirmed as significant decreases based on quantitative morphology (**Figure 2G**). Relatedly, treatment resulted in decreased levels of EpCAM^+^Aqp3^+^ basal cell counts tracked with flow cytometry (**Figure 2H**). In contrast to down-regulation of the basal cell hyperplasia, we found no significant change in SPC^+^IL-33^+^ alveolar-epithelial type 2 (AT2) cell levels with NuP-4A treatment compared to vehicle control (**Figure 2I,J**), providing evidence of basal-epithelial cell selectivity for NuP-4A treatment effect.

**Fig. 2.**
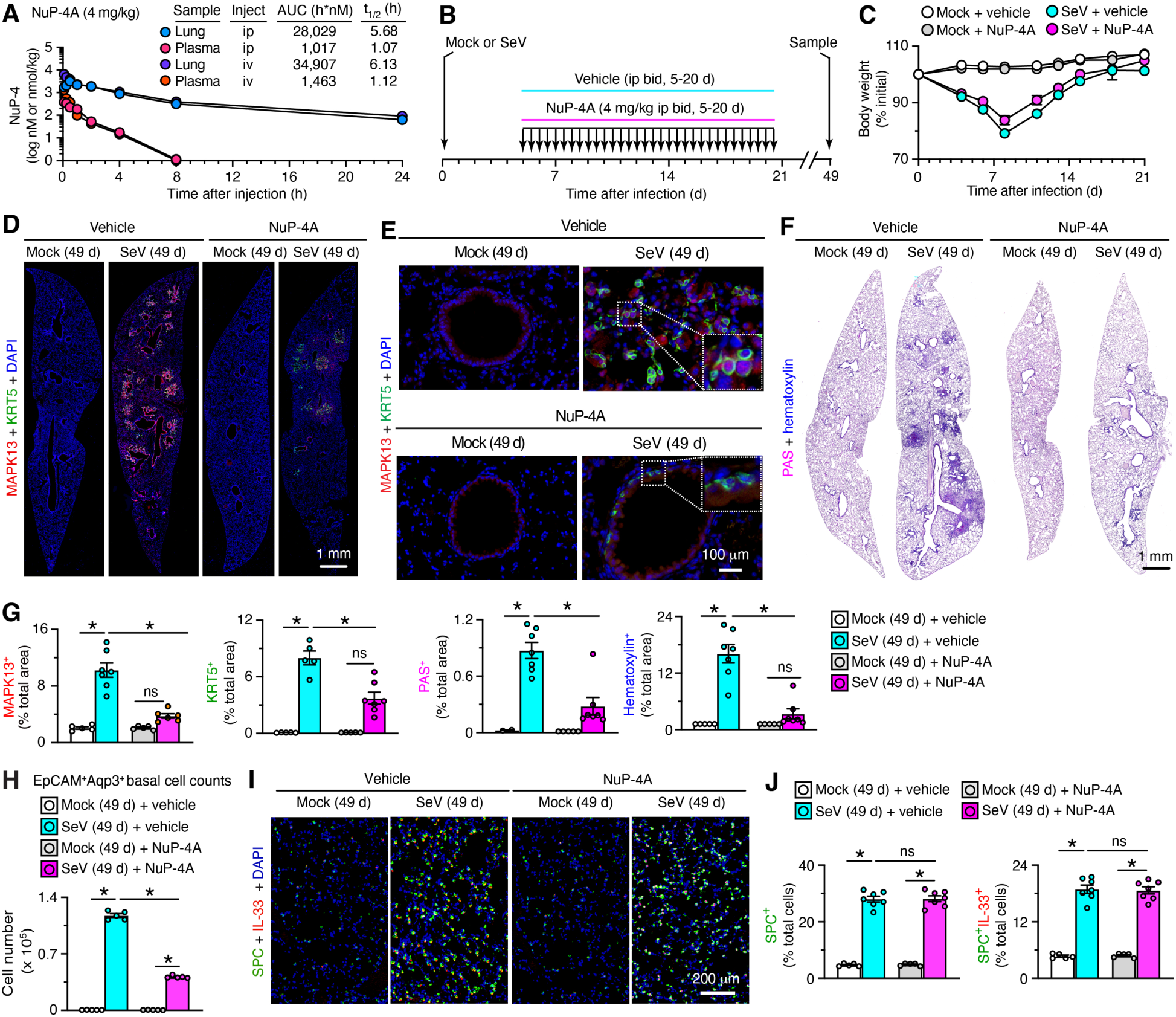
NuP-4A ip treatment attenuates basal-ESC hyperplasia/metaplasia in a mouse model of post-viral asthma. **A**, Levels of NuP-4A (assayed as NuP-4) in lung tissue and plasma after a single dose (4 mg/kg iv or ip) in mice (n=3). AUC, area under curve. **B,** Protocol scheme for NuP-4A or vehicle treatment twice daily (bid) for 5-20 d with final samples at 49 d after SeV infection or Mock control. **C**, Body weights for conditions in (B). **D**, Immunostaining for MAPK13 and KRT5 with DAPI counterstaining in lung sections at 49 d after SeV or Mock and with NuP-4A or vehicle treatment for conditions in (B). E, Higher magnification of immunostaining in (D). Insets are 3x. **F**, PAS-hematoxylin staining of lung sections for conditions in (D). **G**, Quantitation of staining for (D,E). **H**, Cell numbers from flow cytometry of lung epithelial basal cells for conditions in (D). **I,** Immunostaining for SPC and IL-33 with DAPI counterstaining for conditions in (D). **J**, Quantitation of staining for (H). For (C-H), data are representative of three separate experiments with n=5-7 animals per condition in each experiment (mean ± s.e.m.). \**P* <0.05 using ANOVA and Tukey correction.

Treatment with NuP-4 also resulted in significant downregulation of mRNA biomarkers designed to track disease endpoints. This approach showed that treatment caused significant attenuation of biomarkers for basal-ESC hyperplasia marked by *Krt5, Aqp3*, and *Trp63* but not alarmin signal marked by *Il33* (**Figure 3A-E**) consistent with selective MAPK13 control of basal-ESCs but not the larger IL-33^+^ pool of AT2 cells in mouse lungs (19). Treatment also down-regulated immune activation marked by *Serpinb2*, *Ltf, Cxcl17, and Nos2*; type 2 inflammation marked by *Il13, Arg1, Trem2* and *Spp1*; and type1/2 inflammation marked by *Il6* (**Figure 3A-E**) also consistent with MAPK13 control found in *Mapk13*^-/-^ mice (19). No significant treatment effects were found for conventional type 1 inflammation marked by *Ifng, Tnfa,* and *Il1b* mRNA (**Figure 3F**) consistent with data that this signal is often MAPK14-dependent and is not driving chronic disease as found in the present model (19). Relatedly, treatment did not significantly affect the selective induction of *Mapk13* (**Figure 3G**) consistent with relatively small induction and dilution of *Mapk13*^+^ basal-ESC signal. The most potent effect of treatment was found for mucous metaplasia marked by *Muc5ac* and *Clca1* (**Figure 3H**). The *Muc5b* mRNA signal for mucous metaplasia was relatively low (**Figure 3H**), but this finding might be due to probe sensitivity or post-transcriptional regulation given higher signal found by immunostaining (see next section). Treatment resulted in reciprocally lower levels of club cell differentiation marked by *Scgb1a1* mRNA (**Figure 3I**) that is relevant to the basal-ESC shift to mucous cell metaplasia.

**Fig. 3.**
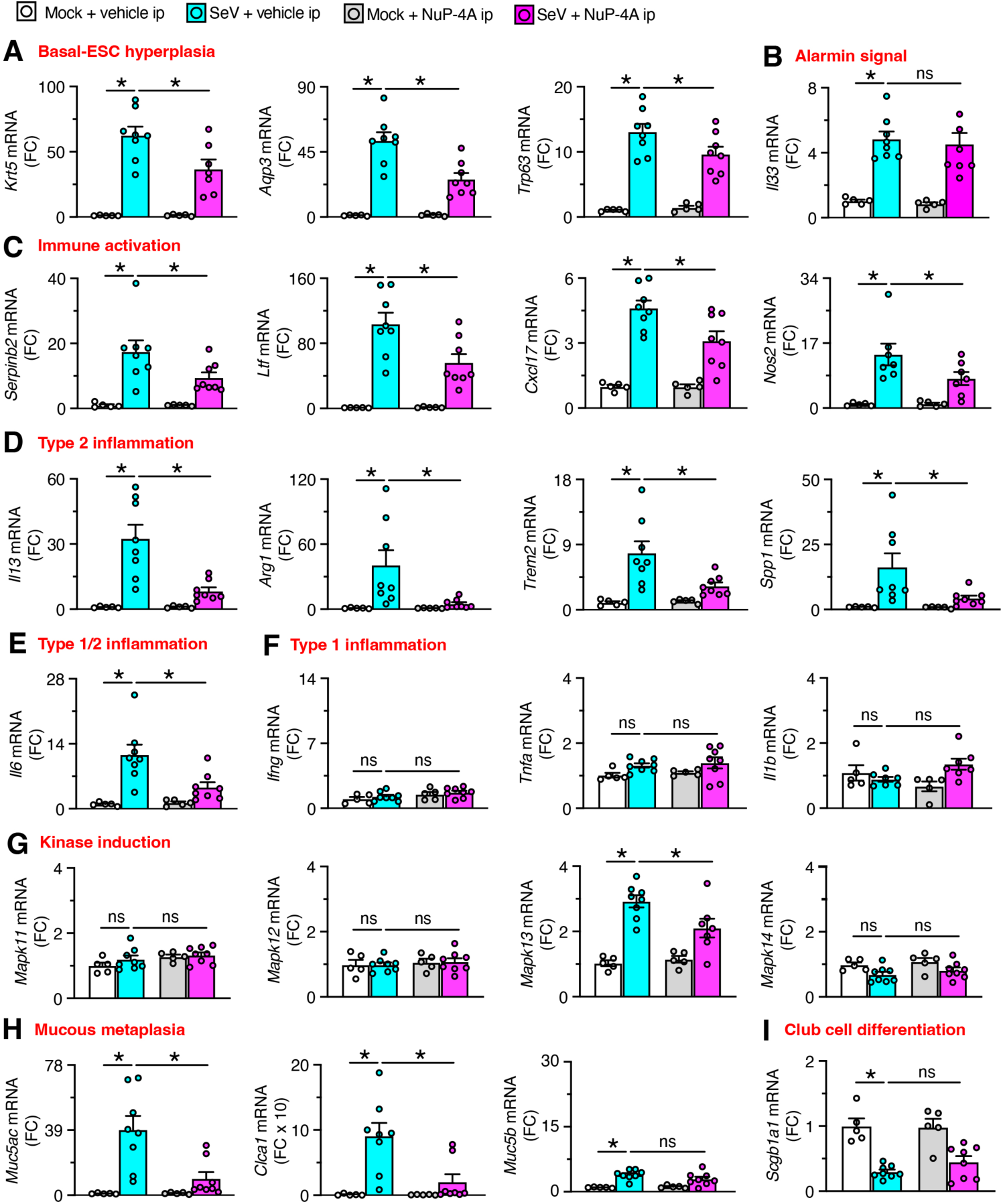
NuP-4A ip treatment attenuates disease biomarkers in a model of post-viral asthma. **A-I**, Lung levels of mRNA biomarkers for indicated disease endpoints for NuP-4A or vehicle treatment twice daily (bid) for 5-20 d with final samples at 49 d after SeV or Mock as shown in Figure 2B. Data are taken from two separate experiments and are representative of three separate experiments with n=5-8 animals per condition in each experiment (mean ± s.e.m.). \**P* <0.05 using ANOVA and Tukey correction.

In concert with mRNA biomarker levels, we also found correction of clinical disease readouts. These included: immune activation based on NOS2^+^ immunostaining (**Figure 4A,B**); macrophage infiltration based on F4/80^+^ immunostaining (**Figure 4C,D**); eosinophil infiltration and degranulation based on major basic protein^+^ (MBP^+^) immunostaining in cellular and extracellular sites (**Figure 4E,F**); mucous metaplasia based on MUC5AC and MUC5B immunostaining (**Figure 4G,H**); and hypoxemia based on SpO_2_ (**Figure 4I**). Notably, lung disease readouts did not recur after stopping treatment, suggesting that proper control of MAPK13 might confer a disease-modifying benefit.

**Fig. 4.**
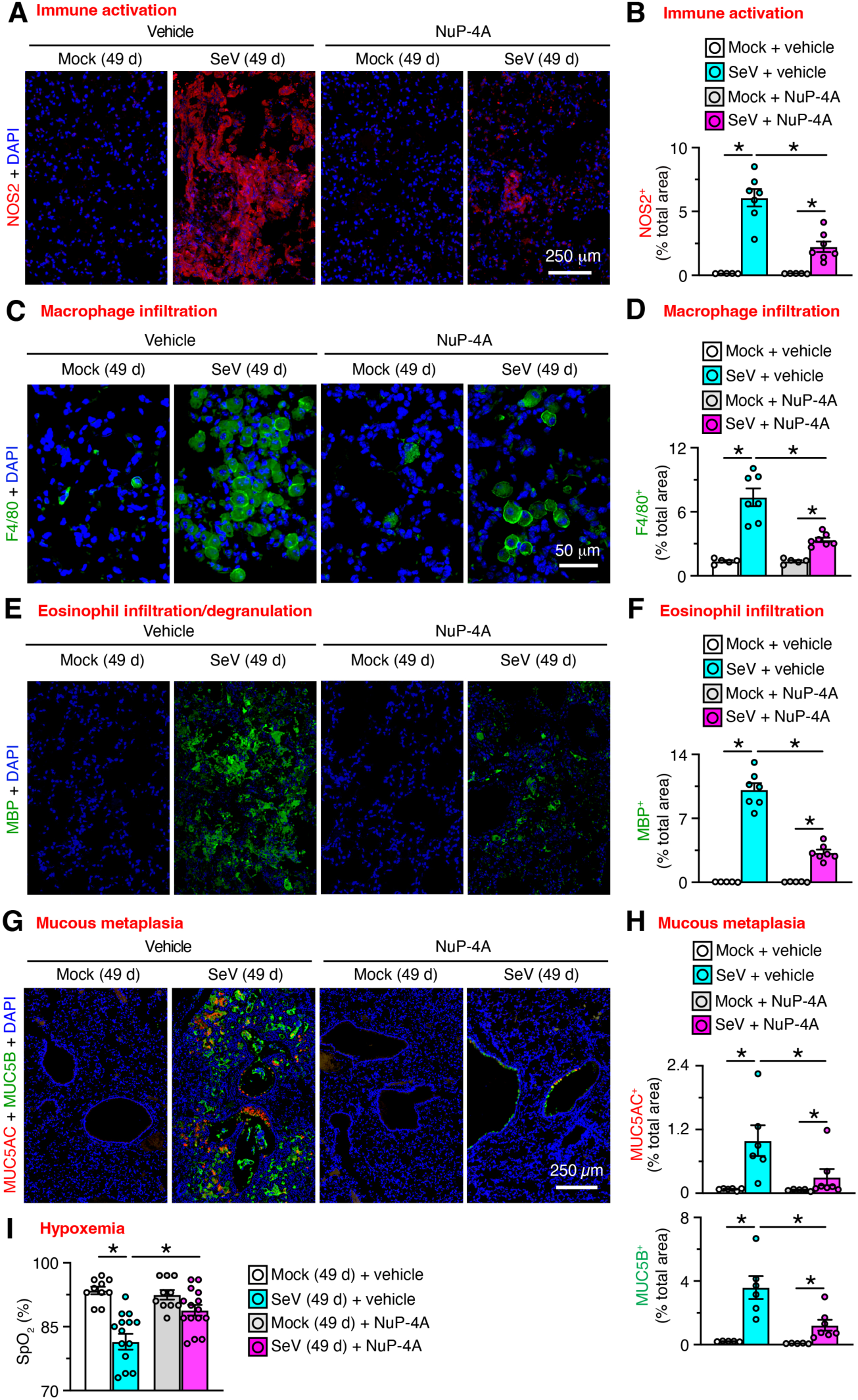
NuP-4A ip treatment attenuates clinical readouts in a model of post-viral asthma. **A**, Immunostaining for NOS2 with DAPI counterstaining of lung sections from mice for NuP-4A treatment for 5-20 d with final samples at 49 d after SeV or Mock as shown in Figure 2B. **B**, Quantitation of staining in (A). **C,** Immunostaining for F4/80^+^ macrophages and DAPI counterstaining for conditions in (A). **D**, Quantitation of staining in (C). **E**, Immunostaining for MBP^+^ eosinophils and DAPI counterstaining for conditions in (A). **F**, Quantitation of staining for (E). **G**, Immunostaining for MUC5AC and MUC5B with DAPI counterstaining for conditions in (A). **H**, Quantitation of staining in (G). **I**, Oximeter levels for blood oxygen saturation (SpO_2_) for conditions in (A). Data are representative of three separate experiments with n=5-14 animals per condition in each experiment (mean ± s.e.m.). \**P* <0.05 using ANOVA and Tukey correction.

We also determined whether NuP-4A was safe and effective even when delivered ahead of infectious illness and was comparable to broad kinase blockade and MAPK13 deficiency. For this approach, NuP-4A treatment was started 2 d before infection and was compared to a combined MAPK13-14 inhibitor NuP-3A and *Mapk13^-/-^*mice as described previously (19, 23). Under this protocol, acute weight loss was attenuated while lung levels for viral RNA and titer were not significantly different among NuP-4A-treated, NuP-3A-treated, *Mapk13^-/-^* mice, and control vehicle-treated mice (**Supplemental Figure 2A-D**). In addition, NuP-4A treatment was as effective as NuP-3A or *Mapk13*-deficiency in attenuating basal cell hyperplasia, mucous metaplasia, and cellularity in lung sections (**Supplemental Figure 2E-G**). Similarly, mRNA biomarkers showed down-regulation of basal-ESC hyperplasia, immune activation, type 2 inflammation, *Mapk13* induction, and mucous metaplasia with no significant change in alarmin signal, type 1 inflammation, or *Muc5b* induction, (**Supplemental Figure 3A-H**). We observed no adverse effects with NuP-4A, thereby providing initial evidence of compound safety under baseline and infection conditions. The findings were similar across interventions (*Mapk13* gene-knockout and both drug treatments) and lung disease readouts at 49-d after infection (**Figs. 2,3**). Together, the data further suggest that NuP-4A benefit is based on down-regulating MAPK13 function and in turn disease readouts.

### MAPK13 inhibitor NuP-4B dpi reverses post-viral lung disease

Given the advantages for drug delivery directly to the lung epithelium, we next tested NuP-4B dpi treatment effect on muco-obstructive lung disease after viral infection. Pharmacokinetic (PK) analysis indicated adequate lung levels were provided with an 80-fold lower dose (achieved with lactose blending) compared to initial dpi dosing as neat compound (**Figure 5A versus Figure 1I**) and a duration of lung levels that was again suitable for once-daily dosing. With this approach, we tested NuP-4B dpi and NuP-4A ip once-daily treatment at 33-48 d after infection that was designed to achieve reversal of established disease (as diagrammed in **Figure 5B**). Under these conditions, NuP-4 treatments showed no adverse effects as marked by stable body weights (**Figure 5C**). Most notably, both NuP-4A ip and NuP-4B dpi treatments achieved marked and equivalent attenuation of the usual increases in MAPK13^+^ cells and KRT5^+^ basal-epithelial cells (**Figure 5D**) along with decreases in corresponding histopathology tracked with PAS and hematoxylin staining (**Figure 5E**). Each of the NuP-4A ip or NuP-4B dpi treatment levels for MAPK13, KRT5, PAS, and hematoxylin staining were significantly decreased compared to vehicle control treatment based on quantitative morphology (**Figure 5F**). In addition, NuP-4A ip or NuP-4B dpi treatment resulted in significantly decreased immune cell infiltration marked by eosinophil and neutrophil levels in bronchoalveolar lavage (BAL) fluid (**Figure 5G**).

**Fig 5.**
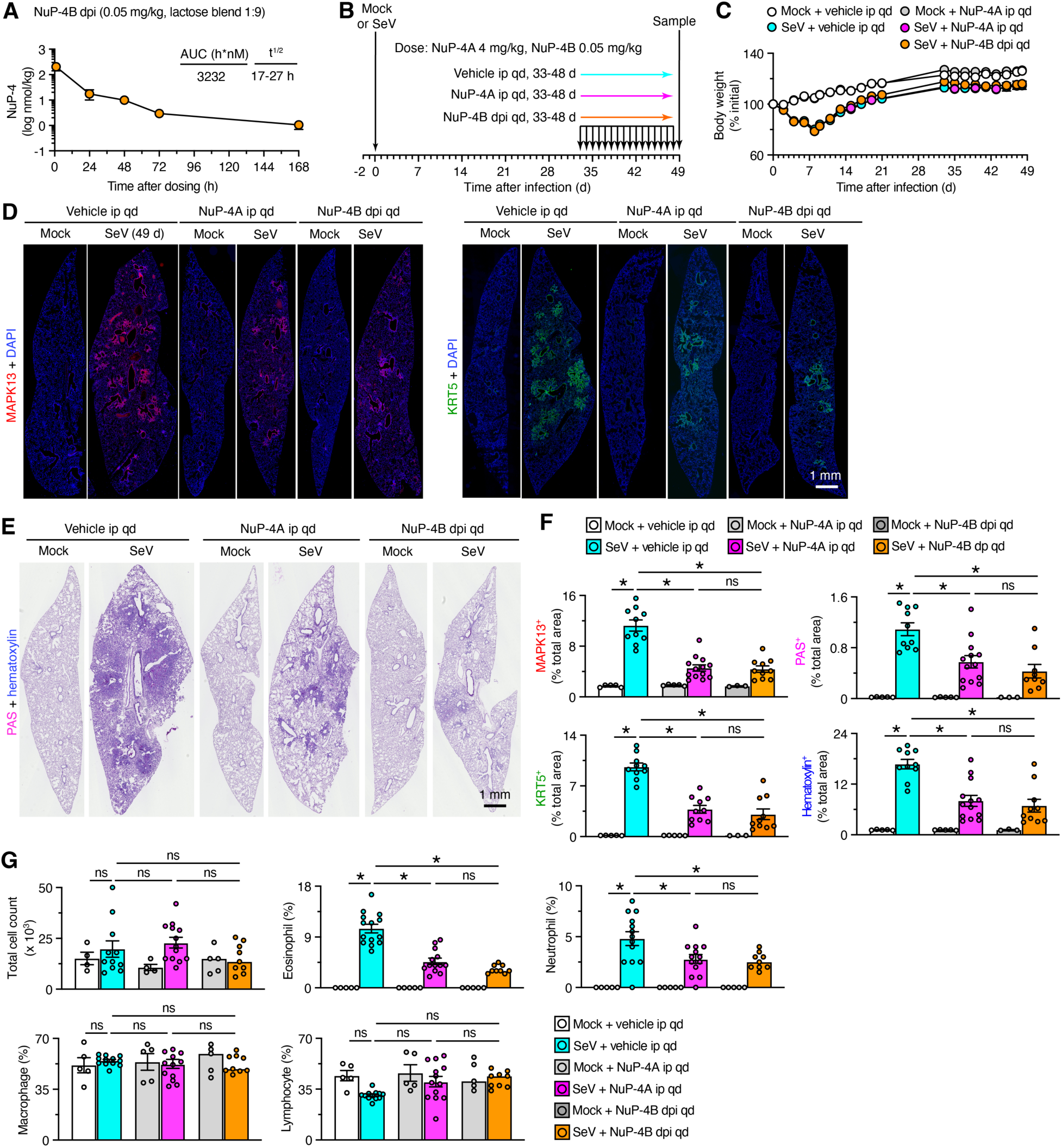
NuP-4A ip or NuP-4B dpi treatment reverses disease in a model of post-viral asthma. **A**, Levels of NuP-4B (assayed as NuP-4) in lung tissue and plasma after a single dose (0.05 mg/kg dpi) in mice (n=3). **B**, Protocol scheme for NuP4A ip, NuP-4B dpi, or vehicle treatment for 33-48 d with samples at 49 d after SeV infection or Mock control. **C**, Body weights for conditions in (B). **D**, Immunostaining for MAPK13 or KRT5 with DAPI counterstaining in lung sections at 49 d after SeV or Mock for conditions in (B). **E**, PAS-hematoxylin staining of lung sections for conditions in (D). **F**, Quantitation of staining for (D,E). **G,** Immune cell counts and percentages in BAL fluid for conditions in (B). Data represent an experiment with n=5-12 animals per condition (mean ± s.e.m.). \**P* <0.05 using ANOVA and Tukey correction.

Treatment with NuP-4A ip or NuP-4B dpi also resulted in significant and similar downregulation of mRNA biomarkers and clinical disease readouts typical of muco-obstructive lung disease. Thus, both forms of NuP-4 treatment attenuated mRNA biomarkers of basal-ESC hyperplasia, immune activation, type 2 inflammation, and mucous metaplasia but not alarmin signal or *Mapk13* induction (**Figure 6A-H**). Relatedly, these treatments significantly down-regulated immunostaining for immune activation marked by NOS2, macrophage infiltration marked by F4/80, and mucous metaplasia marked by MUC5AC and CLCA1 (**Figure 7A-D**). These findings indicated that NuP-4B dpi (delivered at 80-fold lower dose than NuP-4A ip) can still similarly and effectively reverse key aspects of muco-obstructive lung disease, i.e., basal-ESC reprogramming, inflammation, and mucus production, as a guideline for practical application in human lung disease.

**Fig 6.**
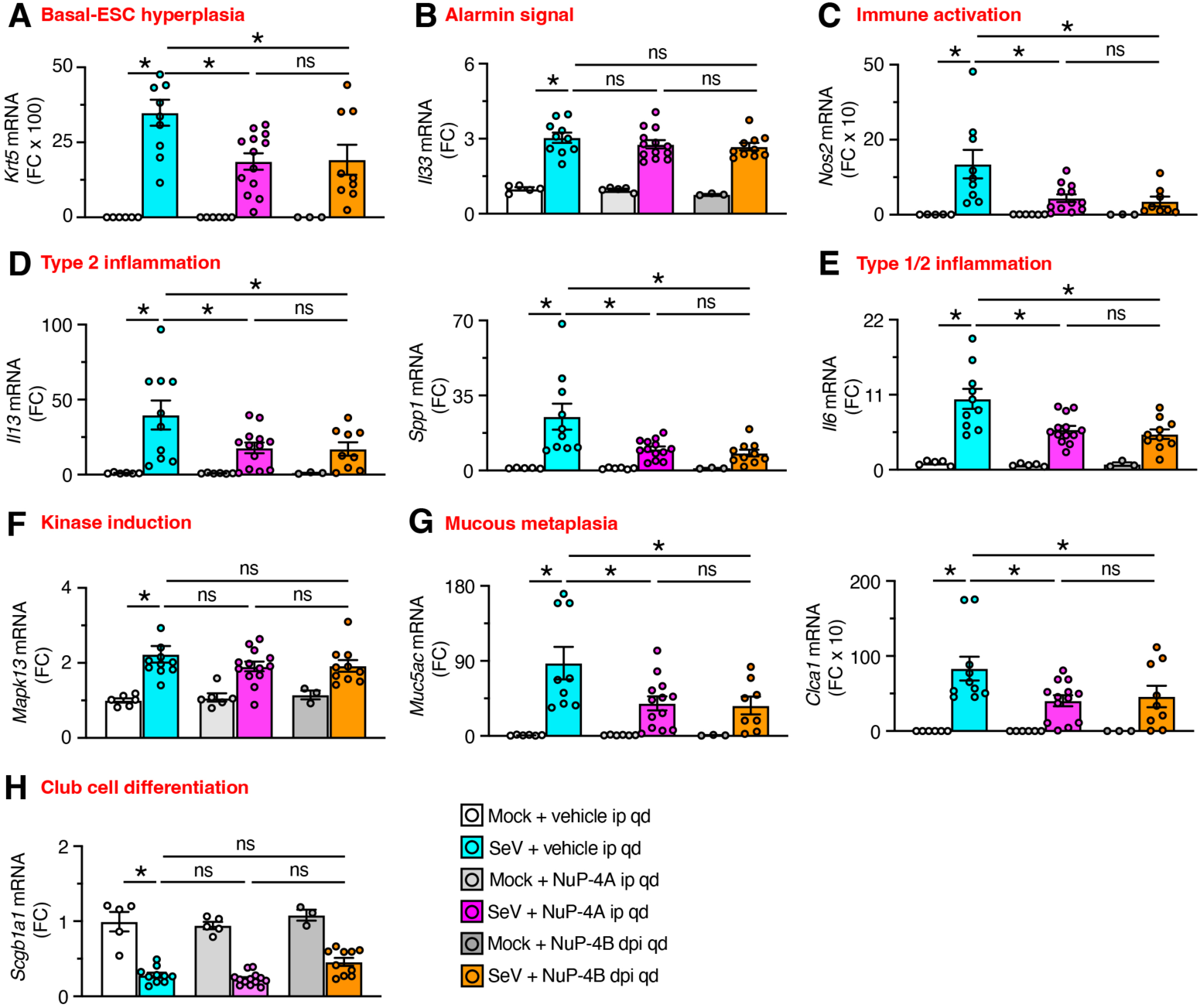
NuP-4A ip or NuP-4B dpi reverses disease biomarkers in a model of post-viral asthma. **A-H**, Lung levels of mRNA biomarkers for indicated disease endpoints for conditions in Figure 5B. Data represent an experiment with n=5-12 animals per condition (mean ± s.e.m.). \**P* <0.05 using ANOVA and Tukey correction.

**Fig 7.**
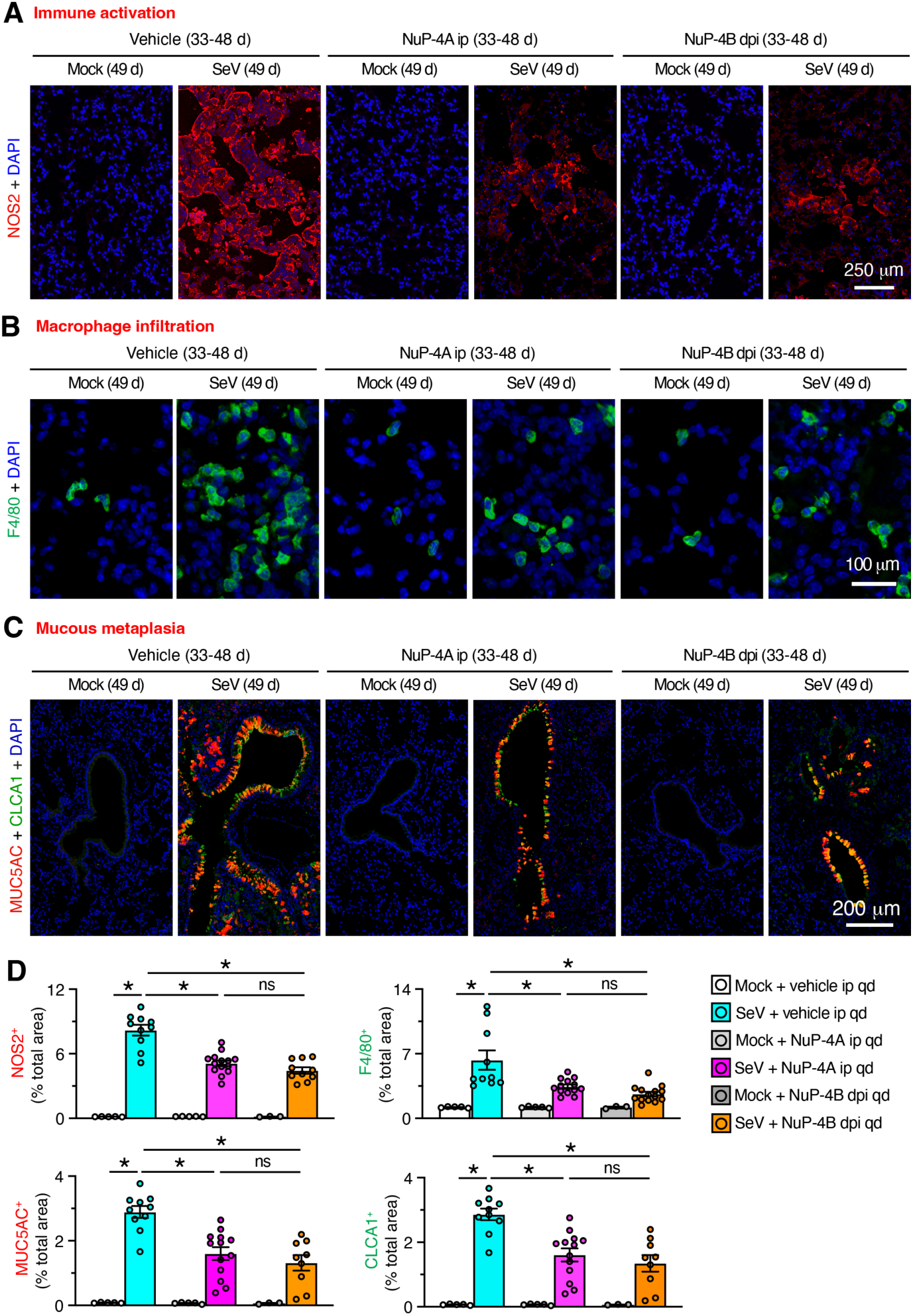
NuP-4A ip or NuP-4B dpi reverses clinical readouts in a model of post-viral asthma. **A-C**, Immunostaining of NOS2, F4/80 or MUC5AC/CLCA1 with DAPI counterstaining in lung sections from conditions in Figure 5B. **D**, Quantitation of staining in (A-C). Data represent an experiment with n=5-12 animals per condition (mean ± s.e.m.). \**P* <0.05 using ANOVA and Tukey correction.

### MAPK13 and MAPK13 inhibitor NuP-4 directly control basal-ESC function

To further extend our experimental findings to practice, we assessed whether MAPK13 expression was found in humans with muco-obstructive lung disease, particularly asthma and COPD that are linked to basal-ESC reprogramming, inflammation, and excess mucus production (18, 33–35). To validate this approach, we characterized lung tissue samples from asthma and COPD patients as described previously (18, 34, 36). The clinical characteristics of lung tissue donors are summarized in **Supplemental Table 1,** recognizing that full features of the medical history are unavailable for some donors that provided tissues collected post-mortem. With this approach, immunostaining of lung tissue sections showed expression of MAPK13 in KRT5^+^ basal-epithelial cells that was detected at sites of mucous cell metaplasia in asthma and COPD (**Figure 8A**). MAPK13 expression was often further localized to a nuclear site, consistent with phosphorylation/activation and nuclear translocation that is typical of MAPK and kinase signaling in general (37). These findings were similar to our model of post-viral lung disease studied here and a related report of this model and human subjects (19).

**Fig. 8.**
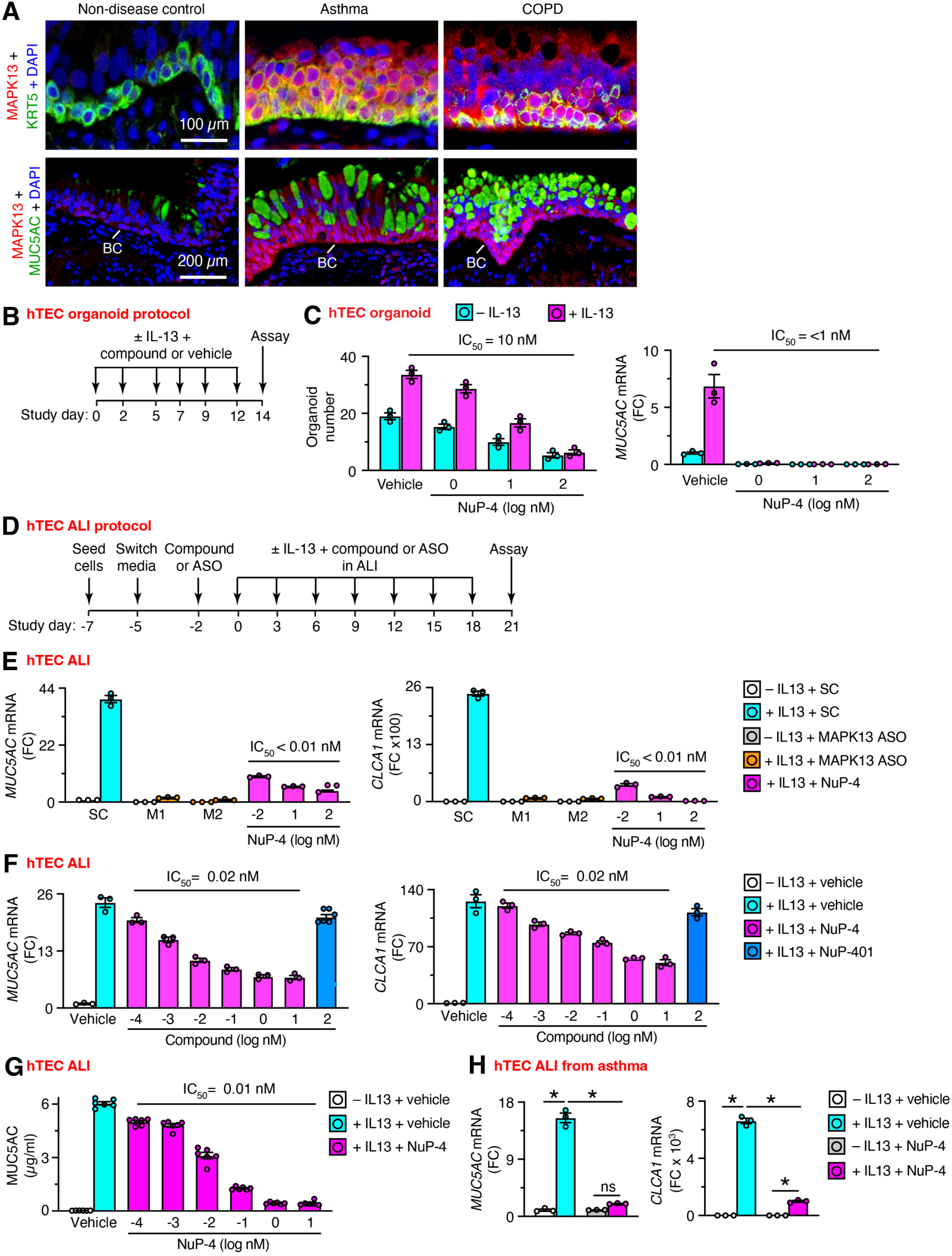
MAPK13 activation and NuP-4 control of human basal-ESCs. **A,** Representative immunostaining for MAPK13 plus KRT5 or MUC5AC with DAPI counterstaining in lung sections from non-disease control (n=11), asthma (n=10), and COPD (n=10) subjects. **B**, Protocol scheme for organoids derived from human tracheobronchial epithelial cells (hTECs) and treated with or without IL-13 and NuP-4 or vehicle. **C**, Levels of cell stemness (marked by organoid formation) and mucous metaplasia (marked by *MUC5AC* mRNA) for conditions in (B). **D**, Protocol scheme for hTECs air-liquid interface (ALI) conditions with or without IL-13, anti-sense oligonucleotide (ASO), and compound or vehicle. **E,** Levels of *MUC5AC* and *CLCA1* mRNA for *MAPK13* ASOs M1 and M2, scrambled control (SC), or NuP-4 treatment using conditions in (D). **F,** Levels of *MUC5AC* and *CLCA1* mRNA for NuP-4 dose-response and maximal-dose NuP-401 using conditions in (D)**. G**, Levels of MUC5AC protein in cell supernatants for conditions in (F). **H**, Levels of *MUC5AC* and *CLCA1* mRNA for conditions in (F) with NuP-4 (100 nM) and cells from an asthma subject. Values (mean ± s.e.m.) derived from a single subject representative of 3 non-disease subjects (C,E,F,G) or 2 asthma subjects (H).

To extend our analysis to MAPK13 and MAPK13-inhibitor control of human basal-ESCs, we developed two new approach methodologies performed ex vivo. In the first model (diagrammed in **Figure 8B**), we used airway organoids derived from primary human tracheobronchial epithelial cells (hTECs) with a new approach methodology modified from our previous report (34). Under these conditions, NuP-4 treatment inhibited baseline and IL-13-enhanced stemness (marked by organoid number) and even more effectively blocked mucous metaplasia (marked by *MUC5AC* mRNA) (**Figure 8C**). Drug effectiveness for stemness (IC_50_ = 10 nM) was equivalent to enzyme-based MAPK13 inhibition based on substrate phosphorylation (**Figure 1C**) but still >10-fold below screening cell toxicity (**Figure 1B**). Notably, NuP-4 effect on mucous metaplasia (IC_50_ <1 nM) was closer to MAPK13-NuP-4 binding affinity (K_D_=20 pM) and thereby >100-fold below screening cell toxicity (**Figure 1E**).

In the second model (diagrammed in **Figure 8D**), we used hTEC-derived cultures under air-liquid interface (ALI) conditions that (when confluent) prevent further growth but still permit IL-13-stimulated differentiation into mucous cells as described previously (18, 23). In this model, we tested treatments with *MAPK13* anti-sense oligonucleotides (ASOs) that decrease MAPK13 levels and with NuP-4 that blocks MAPK13 activity in enzyme and cell-based assays (**Supplemental Figure 4A-F**). Here again, we found that both treatment approaches delivered potent blockade of mucous metaplasia marked with *MUC5AC* and *CLCA1* mRNA levels (**Figure 8E**). Further, dose-response for NuP-4 treatment more precisely defined inhibition of mucous metaplasia at levels remarkably similar to MAPK13–NuP-4 binding affinity (**Figure 8F**). This blocking effect of NuP-4 treatment was not found with a potent and selective MAPK14 inhibitor NuP-401 (VX-745) (38) (**Figure 8F**). Similarly, NuP-4 treatment was also highly effective in down-regulating mucus production marked by secreted MUC5AC level in cell supernatants (**Figure 8G**). Thus, in each of these mucus blocking readouts, NuP-4 efficacy (IC_50_ ≤ 0.02 nM) was nearly identical to binding-kinetics (K_D_=0.02 nM) for NuP-4 that is specific to MAPK13 (**Figure 1E**). For comparison, cell toxicity was not detectable even at 10^6^-fold higher levels of NuP-4 (20 µM) in ALI culture conditions (**Supplemental Figure 4G,H**). For further relevance to lung disease in humans, we also found that NuP-4 treatment delivered similar inhibition of IL-13-stimulated mucous metaplasia in ALI cultures derived from asthma subjects (**Figure 8H**). Together, these results provide evidence that MAPK13 is overexpressed, activated, and susceptible to therapeutic blockade in humans as predicted by our animal models of muco-obstructive lung disease after viral infection.

## Discussion

In this study, we engage drug discovery and proof-of-concept to identify and correct MAPK13 activation as a critical switch-point for basal-ESC reprogramming and muco-obstructive lung disease that develops after viral infection. Key findings include: (1) discovery of the most (and perhaps the first) highly selective MAPK13 inhibitor to date; (2) demonstration that the lead drug candidate (NuP-4) corrects these disease phenotypes similarly to *Mapk13* gene knockout in a mouse model of muco-obstructive lung disease after viral infection; (3) attenuation of disease linked to down-regulation of basal-ESC reprogramming and in turn airway inflammation and mucus production in this model; and (4) corresponding MAPK13 induction/activation in basal-epithelial cells in clinical samples from asthma and COPD patients; and (5) MAPK13 and NuP-4 control of human basal-ESC endpoints (hyperplasia/metaplasia) in new approach methodologies for airway organoid and ALI culture. Together, the data provide a major advance for defining the pathogenesis of chronic lung disease and a practical strategy for modifying this type of disease (as diagrammed in **Figure 9**) and perhaps other diseases proposed for a similar MAPK13-dependent mechanism. Here we highlight three aspects of this research progress.

**Fig. 9.**
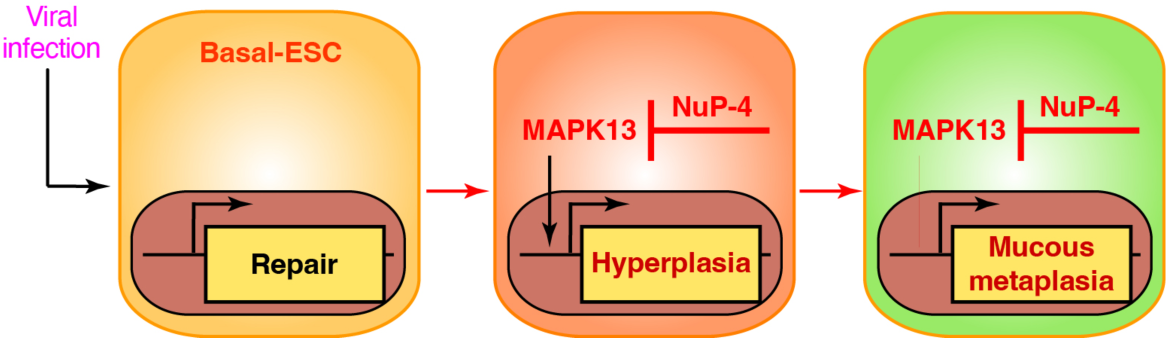
Scheme for MAPK13 inhibitor control of basal-ESC reprogramming as found in post-viral asthma. Sequential steps include viral infection, early homeostatic repair, and later basal-ESC hyperplasia and mucous metaplasia. These long-term endpoints are susceptible to MAPK13 blockade with a small-molecule kinase inhibitor as found in NuP-4. Modified from reference (19).

First, we place the present findings in the context of previous work on the pathogenesis and therapy of chronic lung disease. In that regard, we highlight our studies that defined the role of basal-ESC reprogramming in tissue remodeling and long-term disease after viral infection in the present mouse model (34, 39) and the validation of these findings in humans with chronic lung diseases linked to viral infection, particularly asthma and COPD (34, 35). In parallel with these studies, we identified MAPK13 activation in phospho-kinome screening that translated to studies of *MAPK13* gene-knockdown and blockade of mucous metaplasia in human basal-ESC cultures (18). These initial studies also used a relatively weak MAPK13 inhibitor (parent compound NuP-43; BIRB-796) to demonstrate practical effectiveness. Subsequent drug design based on MAPK13 structure led to first-generation compounds that improved MAPK13 enzyme inhibition but still maintained broad kinase activity (including MAPK14 blockade) (18). Recently, this work was extended to a second-generation MAPK13-14 inhibitor (NuP-3) that was highly effective in blocking mucous metaplasia in human basal-ESC cultures and prevented airway inflammation and mucus production in minipig models using viral infection and cytokine challenge (23). However, this model manifested short-term inflammation and mucus production without the full features of long-term basal-ESC reprogramming and tissue remodeling found in mouse models and humans (34, 35, 39–41). The present work advances these previous approaches in two ways: first, the finding that MAPK13 alone can be sufficient and thereby advantageous as a therapeutic endpoint (based on *MAPK13* gene-knockout in vivo and knockdown in organoid and cell culture); and second, that a more selective MAPK13 inhibitor is achievable and can be sufficient to correct basal-ESC reprogramming and muco-obstructive lung disease (based on treatment effectiveness in animal and human models). Previous work led to the present focus on MAPK13 inhibition, but the present development of a highly selective MAPK13 inhibitor was critical to proof-of-concept and practical guidance.

Second, and in relation to this point, we recognized that it has been historically difficult to effectively and selectively drug MAPK13. Our group was able to take advantage of structural data to reach this endpoint, in contrast to previous approaches often based on kinase screening (42–44). Based on structural modifications of hinge-binding, ATP-binding, and allosteric pocket sites, we achieved significant increases in MAPK13 inhibitor potency, selectivity, binding characteristics, and pharmacokinetic (PK) properties of the lead drug candidate. These advantages are also likely to translate to a decrease in adverse effects in studies of safety. Further, the present development of a formulation for inhaled delivery to the lung epithelium provides additional advantages for effectiveness, selectivity, and consequent safety for properly designed MAPK13 inhibitors and their use in muco-obstructive lung diseases, most commonly in the form of asthma and COPD. In addition, this advance offers clinical guidance to related diseases wherein MAPK13 control of epithelial and immune cells is implicated, but proof-of-concept is based primarily on gene-knockout strategies (14, 45–47). Further, we expect the present advance to provide a tool to bolster the likelihood that MAPK13 maintains a master regulatory role at the epithelial barrier in the lung and other sites. This represents a significant shift from the conventional focus on MAPK14 inhibitors for treatment of inflammation in the lung and other tissue sites (3–6, 8, 11, 48–53). Thus, MAPK14-specific inhibitors are highly effective in blocking conventional cytokine (e.g., TNF-α and IL-1β) signaling (54). This mechanism predicted therapeutic benefit in a variety of conditions including respiratory disease (55); however, even the most advanced versions of these compounds have not proved effective in clinical trials of patients with lung disease, e.g., COPD (8). The data here suggest a basis for this failure by implicating the additional action of MAPK13 in the disease process. As predicted from our earlier work (18), MAPK13 activation is required for type-2 cytokine (particularly IL-13 and IL-4) signaling as a key driver of inflammation and mucus production. Indeed, this predicted the success for blocking type 2 inflammation in asthma and COPD (56). The present data for NuP-4 attenuation of eosinophil infiltration and degranulation along with downregulation of neutrophil infiltration and IL-6 induction suggests a broader benefit for controlling inflammatory disease than is currently available and is required in non-type 2 inflammation conditions.

Third, and relevant to this point, the present data does not exclude an additional role for MAPK13 function in immune cells that act upstream and downstream of basal-ESCs. In that regard, we provided evidence of basal-ESC interactions with the innate immune cell network, including monocyte-derived dendritic cell, macrophage, innate lymphoid cell type 2 (ILC2), and NKT cell populations (33, 57–59). These epithelial-immune cell collaborations thereby provide sentinel-trigger and feed-forward mechanisms for key disease phenotypes, i.e., basal-cell hyperplasia/metaplasia, inflammation, and excess mucus production, that are found in the present mouse model and related disease in humans. Defining the precise role of epithelial-stem cell and perhaps immune-stem cell populations is a challenging goal of future studies. Conditional knockouts of MAPK13 and related members of the CMGC kinase superfamily are feasible (15, 17, 60), but specific tools for epithelial and immune cell subsets needs greater precision. The present development of selective MAPK13 control over basal-ESCs provides practical and precise guidance to correct stem cell reprogramming and consequent disease in the lung and other sites.

## Materials and Methods

### Compound generation, analysis, and formulation

MAPK inhibitor candidates were developed as a chemical analog series (n=520 compounds) from parent compound NuP-43 (BIRB-796) with modifications based on X-ray crystal structures for MAPK13 (unphosphorylated and phosphorylated forms) and co-crystal structure for the MAPK13-NuP-3 complex (18, 23, 25, 26). The entire series of analogs was subjected to a screening funnel that included assessments of chemical properties, hTEC TEER values for cell toxicity at 100 or 1000 nM, and MAPK13 and MAPK14 enzyme inhibition assays. Individual kinase inhibition assays were performed using the HotSpot assay platform (Reaction Biology, Malvern, PA) as described previously (23, 42) using MAP2K6 (MKK6) as upstream kinase activator and myelin basic protein as downstream kinase substrate. The same approach was used to test NuP-43, NuP-3, and NuP-4 for inhibitory activities in a comprehensive human kinase panel (Reaction Biology, Malvern, PA). Binding kinetics for NuP-4 interaction with MAPK12, 13, and 14 were assessed with biolayer interferometry (BLI) as described previously (23).

Lead drug candidate (NuP-4) was formulated as the HBr salt (designated NuP-4A) to improve solubility for all studies of parenteral administration. For pharmacokinetic (PK) analysis, compound levels were determined in plasma and lung tissue samples from mice at a single dose of 2-8 mg/kg and 4 mg/ml in 20% (wt/vol) hydroxypropyl-β-cyclodextrin (Cdx) given by iv or ip injection. Compound levels were determined using a Shimadzu DGU-20A5R(C) HPLC and LCMS-8060 LC/MS/MS instrument or a Prominence Degasser DGU-20A5T(C) HPLC and an AB Sciex Triple Quad 5500 LC/MS/MS instrument. For mouse model experiments, compound was also dissolved in 2-hydroxypropyl-β-cyclodextrin (20% wt/vol) at a compound concentration of 4 mg/ml and stored at 4 °C for each experiment. For experiments with dpi delivery, NuP-4 was formulated as an optimized salt (designated NuP-4B) and prepared as crystalline material for jet-milling. The micronized particles were aerosolized using a rotating brush generator (RBG 1000 ID, Palas GmbH, Karlsruhe, DE) and delivered to mice using an inhalation tower fitted with individual free-roaming tubes (Buxco-DSI, St. Paul, MN). Compounds were used as the freebase form reconstituted in DMSO vehicle (≤1:1000 vol/vol) for cell model experiments. For each preparation, compound stability and purity were verified at >99% using an Agilent 1100 Series HPLC and LC/MSD system and a Varian 400-MHz NMR spectrometer.

### Rat toxicology and toxicokinetic analysis

For rat toxicology studies, NuP-4 was tested in maximum tolerated dose (MTD) and dose-range finding (DRF) protocols for iv dosing using tail vein injection of NuP-4A at 4 mg/ml in 20% 2-hydroxypropyl-β-cyclodextrin solution. Maximal iv doses were limited by feasibility based on NuP-4A solubility and injection volume. Similarly, NuP-4B dpi was delivered to rats as described previously (61) again with dose limited by feasibility at 7 mg/kg for 1 d or 2 mg/kg for 7 d. For both dosing methods, testing was performed in the Sprague-Dawley rat strain, and levels of NuP-4 for lung tissue and plasma samples were determined using a bioanalytical assay with LC/MS/MS detection of NuP-4 as described above.

### Mouse models

Male and female wild-type (WT) C57BL/6J mice (Jax strain 000664) mice were obtained from Jackson Laboratory. The *Mapk13* gene knockout mice (*Mapk13^−/−^*) were generated in the C57BL/6J background using CRISPR/Cas9 technology as described in our preliminary report (19). In brief, with gRNAs that were designed to target early in exon 1 based on a sequence to minimize any off-target effect. Synthetic gRNAs were purchased from IDT Technologies and complexed with recombinant Cas9 protein before nucleofection in Neuro 2a cells for validation. A gRNA that cleaved 30 aa downstream of the start codon of the *Mapk13* gene was chosen to generate the knockout allele. The ribonucleoprotein complex containing Cas9 protein (1 µg/µl) and gRNA (0.3 µg/µl) was then electroporated into single-cell embryos of C57BL/6J mice, and 25–30 eggs were transferred into each pseudo-pregnant female to generate founder mice. Knockout founders were identified by analyzing the PCR-amplified target region (forward primer 5′-ggaacgtacctgggcgag and reverse primer 5′-gtcccacgaactccgagatc) using next-generation sequencing to identify out-of-frame indels. Heterozygous F1 mice were obtained from matings between founders and WT mice, and homozygous F2 mice were obtained through sibling matings of F1 mice. The resulting *Mapk13*^−/−^ mice were found to reproduce and develop normally relative to WT control mice.

All mice were maintained and co-housed in a barrier facility using cages fitted with micro-isolator lids. Animal husbandry and experimental procedures were approved by the Animal Studies Committees of Washington University School of Medicine in accordance with the guidelines from the National Institutes of Health. Sendai virus (SeV, Sendai/52 Fushimi strain, ATCC VR-105) was obtained from ATCC and prepared and titered by plaque-forming assay and qPCR assay as described previously (33). Mice were infected with SeV (2.6 x 10^5^ PFU) as described previously (41). Virus or an equivalent amount of UV-inactivated virus or PBS alone (Mock control) was delivered intranasally in 30 µl of PBS under ketamine/xylazine anesthesia at 6-9 weeks of age. Results from male and female mice were pooled since no significant differences were found between sexes as reported initially (62) and confirmed recently (41) and in the present experiments. Viral titers for stock solutions and lung infections were monitored by quantitative PCR (qPCR) assay using primers for SeV-*NP* RNA as defined previously and in **Supplemental Table 2** using *SeV-NP*-expressing plasmids as an internal standard (41)

### Tissue histology and staining

Lung tissue was fixed with 10% formalin, embedded in paraffin, cut into 5-μm sections and adhered to charged slides. Sections were stained with PAS and hematoxylin as described previously (39, 41). For immunostaining, sections were deparaffinized in Fisherbrand® CitriSolv® (Fisher), hydrated, and heat-treated with antigen unmasking solution (Vector Laboratories, Inc). Immunostaining was performed with the commercially available primary antibodies detailed in **Supplemental Table 3** and a rabbit anti-CLCA1 antibody described previously (18). Primary Abs were detected with secondary Abs labeled with Alexa Fluor 488 (ThermoFisher Scientific) or Alexa Fluor 594 (ThermoFisher Scientific) followed by DAPI counterstaining. Slides were imaged by light microscopy using a Leica DM5000 B and by immunofluorescent microscopy using an Olympus BX51, and staining was quantified in whole lung sections using a NanoZoomer S60 slide scanner (Hamamatsu) and ImageJ software as described previously (39, 41).

### Flow cytometry and FACS

Single cell suspensions were generated from minced lung tissue that was subjected to collagenase (Liberase TM Research Grade, Roche), hyaluronidase (Sigma), DNAse I (Sigma), and Dispase II (Roche) digestion for 45 min at 37 °C and then treated with ACK buffer (Lonza) to remove red blood cells. Following FcR blockade, lung cell suspensions were incubated with labeled antibodies and were sorted using a Sony SY3200 Synergy high-speed cell sorter. The following antibodies were used: anti-mouse CD31 (clone MEC 13.3; BD Biosciences), anti-mouse CD45 (clone 30-F11; BD Biosciences), anti-mouse EpCAM (clone G8.8; BioLegend), anti-AQP3 (Abcam) and anti-Ki-67 (clone SolA15, eBiosciences). Anti-AQP3 antibody was labeled using the Zenon antibody labeling kit (Molecular Probes). FACS results were plotted and analyzed using FlowJo software (TreeStar, Becton Dickinson).

### RNA analysis

RNA was purified from homogenized lung tissue using Trizol (Invitrogen) or from isolated cells with the RNeasy mini kit (Qiagen) and was used to generate cDNA with the High-Capacity cDNA Archive kit (Life Technologies). We quantified target mRNA and viral RNA levels using real-time qPCR assay with specific fluorogenic probe-primer combinations and Fast Universal PCR Master Mix systems (Applied Biosystems) with mouse-specific forward and reverse primers and probes as described previously (59) and in **Supplemental Table 4**. For the *Mapk13* assay, primers and probe were designed to amplify WT but not *Mapk13* indel sequence in *Mapk13*^-/-^ mice. All samples were assayed using the 7300HT or QuantStudio 6 Fast Real-Time PCR System and analyzed using Fast System Software (Applied Biosystems). All real-time PCR data was normalized to the level of *Gapdh* mRNA. Values were expressed as fold-change based on the delta-delta Ct method as described previously (63).

### Lung function tests

To test lung function, we assessed blood oxygen saturation (SpO_2_). For SpO_2_ monitoring, we used a MouseSTAT® pulse oximeter (Kent Scientific, Torrington, CT) applied to the paw skin under isoflurane anesthesia as described previously (39). To decrease variability in readings for this procedure, anesthetized mice were placed on the RightTemp warming pad, and the SpO_2_ reading was recorded when the Oxiwave signal was stable.

### Human clinical samples

For asthma, COPD, and non-disease control samples, lung tissue was obtained from our Advanced Lung Disease Tissue Registry that contains whole lung explants harvested from consented patients in our lung transplant program; lungs that were not usable for transplantation from the local Organ Procurement Organization, Mid-America Transplant; and lungs from a tissue procurement service (IIAM, Edison, NJ) as described previously (18, 35, 36, 64). Human studies were conducted with protocols approved by the Washington University (St. Louis, MO) Institutional Review Board and USAMRDC Office of Research Protections. Clinical characteristics of tissue donors are provided in **Supplemental Table 1**. For asthma and non-disease control samples, clinical information was from deceased donors and therefore was often not available.

### Human epithelial cell culture

Human tracheal and bronchial epithelial cells (hTECs) were isolated by enzymatic digestion and studied to monitor cell growth, mucous metaplasia, and mucus secretion. To monitor cell growth, hTECs were cultured under submerged conditions as described previously (34), and growth was monitored using the CyQUANT cell proliferation assay (ThermoFisher). To monitor mucous metaplasia, hTECs were cultured under air-liquid interface conditions as described previously (18). In this case, cells were cultured in 24-well Transwell plates (6.5-mm diameter inserts, 0.4 µm pore size) from Corning (Corning, NY) with 2% NuSerum medium (65) supplemented with Primocin (50 µg/ml, (InvivoGen, San Diego, CA), and retinoic acid (1 × 10^-8^ M, Sigma, St. Louis, MO) with or without human IL-13 (10 ng/ml, Peprotech, Rocky Hill, NJ) under submerged conditions for 7 d and then air-liquid interface conditions for 21 d. Cells were cultured in the presence or absence of inhibitors or vehicle that were added 2 d before addition of IL-13 and were re-added with each medium change/IL-13 treatment (twice per week). Transepithelial electrical resistance (TEER) was monitored for monolayer integrity as described previously (18, 66). For each set of experiments, cells were assessed with compound treatment versus vehicle control. In addition to 2D culture conditions, hTECs were also cultured under 3D-Matrigel conditions to permit organoid formation as described previously (34). For all conditions, compounds were prepared and stored as 10 mM in DMSO stock solution and for each experiment diluted at least 10,000-fold in cell culture media for cell treatment. Compound effect on mucus production was based on target mRNA and protein levels that were determined as described above using real-time qPCR assay and ELISA. Anisomycin stimulation of MAPK2K3/6 was performed as described previously (67), and LDH release for cytotoxicity was performed using a commercial assay (CyQuant LDH assay kit C20301, Thermo-Fisher).

### Immunoblotting

For western blotting, cultured cells were lysed in RIPA buffer (ThermoFisher) supplemented with PhosSTOP (Sigma) and cOmplete PIC (Sigma) for 30 min at 4 °C. Tissue was homogenized in bead tubes and cell lysates were sheared with a 29G needle, and all samples were centrifuged at 14,100 x g at 4 °C. The supernatant was aliquoted and stored at −80 °C until needed. Protein concentration was determined with the bicinchoninic acid (BCA) protein assay kit (ThermoFisher). Lysate (40-50 µg) was loaded into 4-15% Mini-PROTEAN® TGX gels (Bio-Rad) and separated by SDS-PAGE electrophoresis. Proteins were transferred to nitrocellulose membranes using a Trans-Blot® Turbo blot transfer pack (Bio-Rad). Nitrocellulose membranes were blocked with Odyssey blocking buffer (Li-COR Biosciences) and then primary antibodies for MAPK13 (#/AF1519, R&D) or β-actin (#ab8224, Abcam) were incubated overnight at 4 °C in Odyssey blocking buffer with 0.2% Tween-20. Membranes were washed with 0.1% PBS-Tween-20 and then incubated with anti-mouse or anti-rabbit secondary antibodies (Li-COR) in Odyssey blocking buffer with 0.2% Tween-20 for 1 at 25 °C. Membranes were washed with 0.1% PBS-Tween-20, and protein bands were detected using an Odyssey CLx infrared imaging system (LI-COR Biosciences).

For immunoprecipitation and immunoblotting, lung tissue or cultured cell lysates were treated with SDS (1.5% final concentration), boiled for 10 min, and then diluted 15-fold in TBS buffer. Total protein determination was used to aliquot 500 µg of each sample for immunoprecipitation with 50-µl of Dynabead Protein G slurry (Invitrogen) coupled with 5-µg of anti-MAPK13 mAb (#2308S, Cell Signaling). Samples were heated in 2x-SDS buffer for elution, separated by electrophoresis on 4-12% NuPage Bis-Tris gels (ThermoFisher), and then transferred to PVDF membrane with the Turbo-Blot system as described above. Membranes were blotted for total Mapk13 and phospho-MAPK13 by overnight incubation with anti-MAPK13 polyclonal Ab (#AF1519, R&D Systems) or anti-pan-phospho-p38-MAPK mAb (#9216S, Cell Signaling), respectively. Protein bands were detected and quantified using anti-mouse secondary antibody and the Odyssey CLx imaging system as described above.

### Statistical analysis

All data are presented as mean and s.e.m. and are representative of at least three experiments with at least 5 data points per experiment. Analysis of variance (ANOVA) with Tukey correction for multiple comparisons were used to assess statistical significance between means. In all cases, significance threshold was set at *P* < 0.05.

## Supplemental Materials

Supplemental Figures 1-4 and Tables 1-4.

## Acknowledgments

We thank the Pulmonary Morphology Core and Division of Comparative Medicine for technical support and the NuPeak scientific advisory team (Drs. Bruce Montgomery, Veronika Redmann, Matthew Reed, and Theodore Reiss), Lonza-Bend team (led by Dr. Cameron Kadleck), and Lovelace Biomedical team (led by Dr. Philip Kuehl) for expert input on drug development.

## Author contributions

Y.Z. generated *Mapk13*^-/-^ mice and performed mouse and cell experiments; K.W. performed mouse, cell, and human tissue experiments; D.M. performed mouse and cell experiments; C.A.I performed mouse and cell experiments, H.Y-D. performed immunostaining experiments; K.S. performed mouse experiments; H.W. performed cell culture experiments; S.P.K. performed mouse and cell experiments; M.L. performed structural biology experiments; D.Y. performed mouse breeding experiments; J.Y. performed cell culture experiments, J.R.C. synthesized compounds, Z.H. synthesized compounds, E.C.C. reviewed histopathology; K.O.B. analyzed compound data; D.E.B. obtained and registered human samples; S.L.B. generated human epithelial cell samples; A.G.R. designed and synthesized compounds; and M.J.H. directed the project and wrote the manuscript.

## Abbreviations used in this article

basal-ESC: basal-epithelial stem cell
CLCA1: chloride channel accessory 1
COPD: chronic obstructive pulmonary disease
dpi: dry powder inhalation
GPNMB: glycoprotein non-metastatic melanoma protein B
hTEC: human tracheobronchial epithelial cell
MAPK: mitogen-activated protein kinase
MUC5AC: mucin 5AC
MUC5B: mucin 5B
SeV: Sendai virus.

## References

1. Canovas B, and Nebreda AR. Diversity and versatility of p38 kinase signaling in health and disease. Nat Rev Mol Cell Biol. 2021;22:346–66.

2. Cuenda A, and Sanz-Ezquerro JJ. p38gamma and p38delta: From Spectators to Key Physiological Players. Trends in biochemical sciences. 2017;42(6):431–42.

3. Underwood DC, Osborn RR, and Kotzer CJ. SB 239063, a potent p38 MAP kinase inhibitor, reduces inflammatory cytokine production, airways eosinophil infiltration, and persistence. J Pharmacol Exp Ther. 2000;293:281–8.

4. Duan W, Chan JH, McKay K, Crosby JR, Choo HH, Leung BP, Karras JG, and Wong WSF. Inhaled p38alpha mitogen-activated protein kinase antisense oligonucleotide attenuates asthma in mice. Am J Respir Crit Care Med. 2005;171:571–8.

5. Medicherla S, Fitzgerald MF, Spicer D, Woodman P, Ma JY, Kapoun AM, Chakravarthy S, Dugar S, Protter AA, and Higgins LS. p38a-selective mitogen-activated protein kinase inhibitor SD-282 reduces inflammation in a subchronic model of tobacco smoke-induced airway inflammation. J Pharmacol Exp Ther. 2008;324:921–9.

6. Renda T, Baraldo S, Pelaia G, Bazzan E, Turato G, Papi A, Maestrelli P, Maselli R, Vatrella A, Fabbri LM, Zuin R, Marsico SA, and Saetta M. Increased activation of p38 MAPK in COPD. Eur Respir J. 2008;31:62–9.

7. Millan DS, Bunnage ME, Burrows JL, Butcher KJ, Dodd PG, Evans TJ, Fairman DA, Hughes SJ, Kilty IC, Lemaitre A, Lewthwaite RA, Mahnke A, Mathias JP, Philip J, Smith RT, Stefaniak MH, Yeadon M, and Phillips C. Design and synthesis of inhaled p38 inhibitors for the treatment of chronic obstructive pulmonary disease. J Med Chem. 2011;54:7797–814.

8. MacNee W, Allan RJ, Jones I, Cristina De Salvo M, and Tan LF. Efficacy and safety of the oral p38 inhibitor PH-797804 in chronic obstructive pulmonary disease: a randomised clinical trial. Thorax. 2013;68:738–45.

9. Watz H, Barnacle H, Hartley BF, and Chan R. Efficacy and safety of the p38 MAPK inhibitor losmapimod for patients with chronic obstructive pulmonary disease: a randomised, double-blind, placebo-controlled trial. Lancet Respir Med. 2014;2:63–72.

10. Patel NR, Cunoosamy DM, Fageras M, Taib Z, Asimus S, Hegelund-Myrback T, Lundin S, Pardali K, Kurian N, Ersdal E, Kristensson C, Korsback K, Palmer R, Brown MN, Greenaway S, Siew L, Clarke GW, Rennard SI, Make BJ, Wise RA, and Jansson P. The developmen tof AZD7624 for prevention of exacerbations in COPD: a randomized controlled trial. Int J COPD. 2018;13:1009–19.

11. Haller V, Nahidino P, Forster M, and Laufer SA. An updated patent review of p38 MAPK kinase inhibitors (2014-2019). Expert Opin Ther Pat. 2020;30:453–66.

12. Goedert M, Hasegawa M, Jakes R, Lawler S, Cuenda A, and Cohen P. Phosphorylation of microtube-associated protein tau by stress-activated protein kinases. FEBS Lett. 1997;409:57–62.

13. Sumara G, Formentini I, Collins S, Sumara I, Windak R, Bodenmiller B, Ramracheya R, Caille D, Jiang H, Platt KA, Meda P, Aebersold R, Rorsman P, and Ricci R. Regulation of PKD by the MAPK p38δ in insulin secretion and glucose homeostasis. Cell. 2009;136:235–48.

14. Ittner A, Block H, Reichel CA, Varjosalo M, Gehart H, Sumara G, Gstaiger M, Krombach F, Zarbock A, and Ricci R. Regulation of PTEN activity by p38d-PKD1 signaling in neutrophils confers inflammatory responses in the lung. J Exp Med. 2012;209:2229–46.

15. Schindler E, Hindes A, Gribben E, Burns C, Yin Y, Lin M, Owen R, Longmore G, Kissling G, Arthur J, and Efimova T. p38delta Mitogen-activated protein kinase is essential for skin tumor development in mice. Cancer Res. 2009;69:4648–55.

16. Risco A, Del Fresno C, Mambol A, Alsina-Beauchamp D, MacKenzie KF, Yang H-T, Barber DF, Morcelle C, Arthur JSC, Ley SC, Ardavin C, and Cuenda A. p38g and p38d kinases regulate the Toll-like receptor 4 (TLR4)-induced cytokine production by controlling ERK1/2 protein kinase pathway activation. Proc Natl Acad Sci U S A. 2012;109:11200–5.

17. Tomas-Loba A, Manieri E, Gonzalez-Teran B, Mora A, Leiva-Vega L, Santamans AM, Romero-Becerra R, Rodriguez E, Pintor-Chocano A, Feixas F, Lopez JA, Caballero B, Trakala M, Blanco O, Torres JL, Hernandez-Cosido L, Montalvo-Romeral V, Matesanz N, Roche-Molina M, A. BJ, Mischo H, Leon M, Caballero A, Miranda-Saavedra D, Ruiz-Cabello J, Nevzorova YA, Cubero FJ, Bravo J, Vazquez J, Malumbres M, Marcos M, Osuna S, and Sabio G. p38g is essential for cell cycle progression and liver tumoriogenesis. Nature. 2019;568:557–60.

18. Alevy Y, Patel AC, Romero AG, Patel DA, Tucker J, Roswit WT, Miller CA, Heier RF, Byers DE, Brett TJ, and Holtzman MJ. IL-13–induced airway mucus production is attenuated by MAPK13 inhibition. J Clin Invest. 2012;122:4555–68.

19. Wu K, Zhang Y, Mao D, Iberg C, Yin-Declue H, Sun K, Keeler S, Wikfors H, Young D, Yantis J, Austin S, Byers D, Brody S, Crouch E, Romero A, and Holtzman M. MAPK13 controls structural remodeling and disease after epithelial injury. bioRxiv. 2024;10.1101/2024.05.31.596863.

20. Radicioni G, Ceppe A, Ford AA, Alexis NE, Barr RG, Bleecker ER, Christenson SA, Cooper CB, Han MK, Hansel NN, Hastie AT, Hoffman EA, Kanner RE, Martinez FJ, Ozkan E, Paine R, 3rd, Woodruff PG, O’Neal WK, Boucher RC, and Kesimer M. Airway mucin MUC5AC and MUC5B concentrations and the initiation and progression of chronic obstructive pulmonary disease: an analysis of the SPIROMICS cohort. Lancet Respir Med. 2021;9(11):1241–54.

21. Tang M, Elicker BM, Henry T, Gierada DS, Schiebler ML, Huang BK, Peters MC, Castro M, Hoffman EA, Fain SB, Ash SY, Choi JY, Hall C, Phillips BR, Mauger DT, Denlinger LC, Jarjour NN, Israel E, Phipatanakul W, Levy BD, Wenzel SE, Bleecker ER, Woodruff PG, Fahy JV, Dunican EM, and Program-3 NSAR. Mucus plugs persist in asthma, and changes in mucus plugs associate with changes in airflow over time. Am J Respir Crit Care Med. 2022;205:1036–45.

22. Mettler SK, Nardelli P, Campo MI, San Jose Estepar R, Manapragada PP, Abozeed M, Aziz MU, Zahid M, Grumley S, Nath HP, Yen A, Sonavane S, Clarenbach CF, Wang W, Pistenmaa CL, Washko GR, Estepar RSJ, Cho MH, and Diaz AA. Longitudinal Changes in Airway Mucus Plugs and FEV(1) in COPD. N Engl J Med. 2025;392(19):1973–5.

23. Keeler SP, Wu K, Zhang Y, Mao D, Li M, Iberg CA, Austin SP, Glaser SA, Yantis J, Podgorny S, Brody SL, Chartock JR, Han Z, Byers DE, Romero AG, and Holtzman MJ. A potent MAPK13-14 inhibitor prevents airway inflammation and mucus production. Am J Physiol Lung Cell Mol Physiol. 2023;325:L726–40.

24. Pelaia C, Vatre, A., Gallelli L, Lombardo N, Sciacqua A, Savino R, and Pelaia G. Role of p38 mitogen-activated protein kinase in asthma and COPD: pathogenic aspects and potential targeted therapies. Drug Design, Development and Therapy. 2021;15:1275–84.

25. Yurtsever Z, Schaeffer SM, Romero AG, Holtzman MJ, and Brett TJ. The crystal structure of phosphorylated MAPK13 reveals common structural features and diffeerences in p38 MAPK family activation. Acta Crystallogr D Biol Crystallogr. 2015;71:790–9.

26. Yurtsever Z, Patel DA, Kober DL, Su A, Miller CA, Romero AG, Holtzman MJ, and Brett TJ. First comprehensive structure and biophysical analysis of MAPK13 inhibitors targeting DFG-in and DFG-out binding modes. Biochim Biophys Acta. 2016;1860:2335–44.

27. Holtzman MJ, Romero AG, Gerovac BJ, Han Z, Keeler SP, and Wu K. In: USPTO ed. United States Patent Application Publication. US: Washington University; 2023.

28. Pargellis C, Tong L, Churchill L, Cirillo PF, Gilmore T, Graham AG, Grob PM, Hickey ER, Moss N, Pav S, and Regan J. Inhibition of p38 MAP kinase by utilizing a novel allosteric binding site. Nat Struct Biol. 2002;9:268–72.

29. Regan J, Pargellis CA, Cirillo PF, Gilmore T, Hickey ER, Peet GW, Proto A, Swinamer A, and Moss N. The kinetics of binding to p38 MAP kinase by analogues of BIRB 796. Bioorg Med Chem. 2003;13:3101–4.

30. Casasnovas R, Limongelli V, Tiwary P, Carloni P, and Parrinello M. Unbinding kinetics of a p38 MAP kinase type II inhibitor from metadynamics simulations. J Am Chem Soc. 2017;139:4780–8.

31. Huang Y. Multiscale computational study of ligand binding pathways: case of p38 MAP kinase and its inhibitors. Biophysical J. 2021;120:3881–92.

32. Pasqua E, Hamblin N, Edwards C, Baker-Glenn C, and Hurley C. Developing inhaled drugs for respiratory diseases: a medicinal chemistry perspective. Drug Disc Today. 2022;27:134–50.

33. Kim EY, Battaile JT, Patel AC, You Y, Agapov E, Grayson MH, Benoit LA, Byers DE, Alevy Y, Tucker J, Swanson S, Tidwell R, Tyner JW, Morton JD, Castro M, Polineni D, Patterson GA, Schwendener RA, Allard JD, Peltz G, and Holtzman MJ. Persistent activation of an innate immune response translates respiratory viral infection into chronic lung disease. Nat Med. 2008;14:633–40.

34. Byers DE, Alexander-Brett J, Patel AC, Agapov E, Dang-Vu G, Jin X, Wu K, You Y, Alevy YG, Girard J-P, Stappenbeck TS, Patterson GA, Pierce RA, Brody SL, and Holtzman MJ. Long-term IL-33-producing epithelial progenitor cells in chronic obstructive lung disease. J Clin Invest. 2013;123:3967–82.

35. Wu K, Zhang Y, Yin Declue H, Austin SR, Byers DE, Crouch EC, and Holtzman MJ. Lung remodeling regions in long-term coronavirus disease 2019 feature basal epithelial cell reprogramming. Am J Pathol 2023;193:680–9.

36. Deslee G, Woods J, Moore C, Conradi S, Gierada D, Atkinson J, Battaile J, Liu L, Patterson A, Adair-Kirk T, Holtzman M, and Pierce R. Oxidative damage to nucleic acids in severe emphysema. Chest. 2009;135:965–74.

37. Holtzman MJ, Zhang Y, Wu K, and Romero AG. Mitogen-activated protein kinase-guided drug discovery for post-viral and related types of lung disease. Eur Respir Rev. 2024;33:230220.

38. Duffy JP, Harrington EM, Salituro FG, Cochran JE, Green J, Gao H, Bemis GW, Evindar G, Galulio VP, Ford PJ, Germann UA, Wilson KP, Bellon SF, Chen G, Taslimi P, Jones P, Huang C, Pazhanisamy S, Wang YM, Murcko MA, and Su MS. The discovery of VX-745: a novel and selective p38a kinase inhibitor. ACS Med Chem Lett. 2011;2:758–63.

39. Wu K, Kamimoto K, Zhang Y, Yang K, Keeler SP, Gerovac BJ, Agapov EV, Austin SP, Yantis J, Gissy KA, Byers DE, Alexander-Brett J, Hoffmann CM, Wallace M, Hughes ME, Morris SA, and Holtzman MJ. Basal-epithelial stem cells cross an alarmin checkpoint for post-viral lung disease. J Clin Invest. 2021;131:e149336.

40. Keeler SP, Agapov EV, Hinojosa ME, Letvin AN, Wu K, and Holtzman MJ. Influenza A virus infection causes chronic lung disease linked to sites of active viral RNA remnants. J Immunol. 2018;201:2354–68.

41. Zhang Y, Mao D, Keeler SP, Wang X, Wu K, Gerovac BJ, Shornick LP, Agapov E, and Holtzman MJ. Respiratory enterovirus (like parainfluenza virus) can cause chronic lung disease if protection by airway epithelial STAT1 is lost. J Immunol. 2019;202:2332–47.

42. Anastassiadis T, Deacon SW, Devarajan K, Ma H, and Peterson JR. Comprehensive assay of kinase catalytic activity reveals features of kinase inhibitor selectivity. Nat Biotechnol. 2011;29:1039–45.

43. Klaeger S, Heinzlmeir S, Wilhelm M, Polzer H, Vick B, Koenig P-A, Reinecke M, Ruprecht B, Petzoldt S, Meng C, Zecha J, Reiter K, Qiao H, Helm D, Koch H, Schoof M, Canevari G, Casale E, Depaolini SR, Feuchtinger A, Wu Z, Schmidt T, Rueckertr L, Becker W, Huenges J, Garz A-K, Gohlke B-O, Zolg DP, Kayser G, Vooder T, Preissner R, Hahne H, Tonisson N, Kramer K, Gotze K, Bassermann F, Schlegl J, Ehrlich H-C, Aiche S, Walch A, Greif PA, Schneider S, Felder ER, Ruland J, Medard G, Jeremias I, Spiekermann K, and Kuster B. The target landscape of clinical kinase drugs. Science. 2017;358:1148.

44. Cohen P, Cross D, and Janne PA. Kinase drug discovery 20 years after imatinib: progress and future directions. Nat Rev Drug Discov. 2021;20:551–69.

45. Alsina-Beauchamp D, Escos A, Fajardo P, Gonzalez-Romero D, Diaz-Mora E, A. R, Martin-Serrano MA, C. dF, Dominguez-Andres J, Aparicio N, Zur R, Shpiro N, Brown GD, Ardavin C, Netea MC, Alemany S, Sanz-Ezquerro JJ, and Cuenda A. Myeloid cell deficiency of p38g/p38d protects against candidiasis and regulates antifungal immunity. EMBO Mol Med. 2018;10:e8485.

46. Escos A, Martin-Gomez J, Gonzalez-Romero D, Diaz-Mora E, Francisco-Velilla R, Santiago C, Cueza J, Dominguez-Zorita S, Martinez-Salas E, Sonenberg N, Sanz-Ezquerro J, Jafanejad S, and Cuenda A. TPL2 kinase expression is regulated by the p38g/p38d-dependent association with aconitase-1 with *TPL2* mRNA. Proc Natl Acad Sci U S A. 2022;119:e2204752119.

47. Escos A, Diaz-Mora E, Pattison M, Fajardo P, Gonzalez-Romero D, Risco A, Martin-Gomez J, Bonnell E, Sonenberg N, Jafanejad SM, Sanz-Ezquerro JJ, Ley SC, and Cuenda A. p38g and p38d modulate innate immune response by regulating MEF2D activation. eLife. 2023;12:e86200.

48. Kawai T, and Akira S. Pathogen recognition with Toll-like receptors. Curr Opin Immunol. 2005;17(4):338–44.

49. Ha U, H. LJ, Jono H, Koga T, Srivastava A, Malley R, Pages G, Pouyssegur J, and Li J-D. A novel role for IkB kinase (IKK) a and IKKb in ERK-dependent up-regulation of MUC5AC mucin transcription by Streptococcus pneumoniae. J Immunol. 2007;178:1736–47.

50. Fujisawa T, Velichko S, Thai P, Hung LY, Huang F, and Wu R. Regulation of airway MUC5AC expression by IL-1beta and IL-17A; the NF-kappaB paradigm. J Immunol. 2009;183(10):6236–43.

51. Na HG, Kim Y-D, and Bae CH. High concentration of insulin induces MUC5AC expression via phosphoinositide 3 kinase/AKT and mitogen-activated protein kinase signaling pathways in human airway epithelial cells. Am J Rhinol Allergy. 2018.

52. Xu H, Sun Q, Lu L, Luo F, Zhou L, Liu J, Cao L, Wang Q, Xue J, Yang Q, Yang P, Lu J, Xiang Q, and Liu Q. MicroRNA-218 acts by repressing TNFR1-mediated activation of NF-kB, which is involved in MUC5AC hyper-production and inflammation in smoking-induced bronchiolitis of COPD. Toxicol Lett. 2017;280:171–80.

53. Wu S, Li H, Yu L, Wang N, Li X, and Chen W. IL-1b upregulates Muc5ac expression via NF-kB-induced HIF-1a in asthma. Immunol Lett. 2017;192:20–6.

54. Selness SR, Devraj RV, Devadas B, Walker JK, Boehm TL, Durley RC, Shieh H, Xing L, Rucker PV, Jerome KD, and Benson AG. Discovery of PH-797804, a highly selective and potent inhibitor of p38 MAP kinase. Bioorg Med Chem Lett. 2011;21:4066–71.

55. McQualter JL, Yuen K, Williams B, and Bertoncello I. Evidence of an epithelial stem/progenitor cell hierarchy in the adult mouse lung. Proc Natl Acad Sci U S A. 2010;107(4):1414–9.

56. Bhatt SP, Rabe KF, Hanania NA, Vogelmeier CF, Cole J, Bafadhel M, Christenson SA, Papi A, Singh D, Laws E, Mannent LP, Patel N, Staudinger HW, Yancopoulos GD, Mortensen ER, Akinlade B, Maloney J, Lu X, Bauer D, Bansal A, Robinson LB, and Abdulai RM. Dupilumab for COPD with type 2 inflammation indicated by eosinophil counts. N Engl J Med. 2023;389:205–14.

57. Wu K, Byers DE, Jin X, Agapov E, Alexander-Brett J, Patel AC, Cella M, Gilfilan S, Colonna M, Kober DL, Brett TJ, and Holtzman MJ. TREM-2 promotes macrophage survival and lung disease after respiratory viral infection. J Exp Med. 2015;212:681–97.

58. Wang X, Wu K, Keeler SP, Mao D, Agapov EV, Zhang Y, and Holtzman MJ. TLR3-activated monocyte-derived dendritic cells trigger progression from acute viral infection to chronic disease in the lung. J Immunol. 2021;206:1297–314 (selected for Top-Reads p. 115).

59. Wu K, Wang X, Keeler SP, Gerovac BJ, Agapov E, Byers DE, Gilfillan S, Colonna M, Zhang Y, and Holtzman MJ. Group 2 innate lymphoid cells must partner with the myeloid-macrophage lineage for long-term postviral lung disease. J Immunol. 2020;205:1084–101.

60. Varjosalo M, Keskitalo S, Van Drogen A, Nurkkala H, Vichalkovski A, Aebersold R, and Gstaiger M. The protein interaction landscape of the human CMGC kinase group. Cell Rep. 2013;3(1306-1320).

61. Tepper JS, Kuehl PJ, Cracknell S, Nikula KJ, Pei L, and Blanchard JD. Symposium summary: “Breathe in, breathe out, its easy: what you need to know about developing inhaled drugs”. Int J Toxicol. 2016:1–17.

62. van Nunen MCJ, and van der Veen J. Experimental infection with Sendai virus in mice. Arch Gesamte Virusforsch. 1967;22:388–97.

63. Livak KJ, and Schmittgen TD. Analysis of relative gene expression data using real-time quantitative PCR and the 2(-Delta Delta C(T)) Method. Methods. 2001;25:402–8.

64. Byers DE, Wu K, Dang-Vu G, Jin X, Agapov E, Zhang X, Battaile JT, Schechtman KB, Yusen R, Pierce RA, and Holtzman MJ. Triggering receptor expressed on myeloid cells-2 (TREM-2) expression tracks with M2-like macrophage activity and disease severity in COPD. Chest. 2018;153:77–86.

65. Lechner JF, and LaVeck MA. A serum-free method for culturing normal human bronchial epithelial cells at clonal density. J Tissue Culture Meth. 1985;9:43–8.

66. Patel DA, Patel AC, Nolan WC, Zhang Y, and Holtzman MJ. High throughput screening for small molecule enhancers of the interferon signaling pathway to drive next-generation antiviral drug discovery. PloS ONE. 2012;7:e36594.

67. Remy G, Risco A, Inesta-Vaquera F, Gonzalez-Teran B, Sabio G, Davis R, and Cuenda A. Differential activation of p38MAPK isoforms by MKK6 and MKK3. Cellular Signaling. 2010;22:660–7.

