## Supplemental Materials for "A first-in-kind MAPK13 inhibitor corrects epithelial stem cell reprogramming and muco-obstructive lung disease"

#### Supplemental Figures 1-4 and Tables 1-4

### A first-in-kind MAPK13 inhibitor corrects stem cell reprogramming and muco-obstructive lung disease

**Yong Zhang[1*], Kangyun Wu[1*], Dailing Mao[1], Courtney A. Iberg[1], Huiqing Yin-Declue[1], Kelly Sun[1], Hallie A. Wikfors[1], Shamus P. Keeler[1], Ming Li[1], Deanna Young[1], Jennifer Yantis[1], Joshua R. Chartock[1], Zhenfu Han[1], Erika C. Crouch[2], Kay O. Broschat[4], Derek E, Byers[1], Steven L. Brody[1], Arthur G. Romero[1], and Michael J. Holtzman[1,3,4*]**

[1]Pulmonary and Critical Care Medicine, Department of Medicine, [2]Department of Pathology and Immunology, and [3]Department of Cell Biology and Physiology, Washington University School of Medicine, St. Louis, MO 63110 and [4]NuPeak Therapeutics Inc., St. Louis, MO 63105

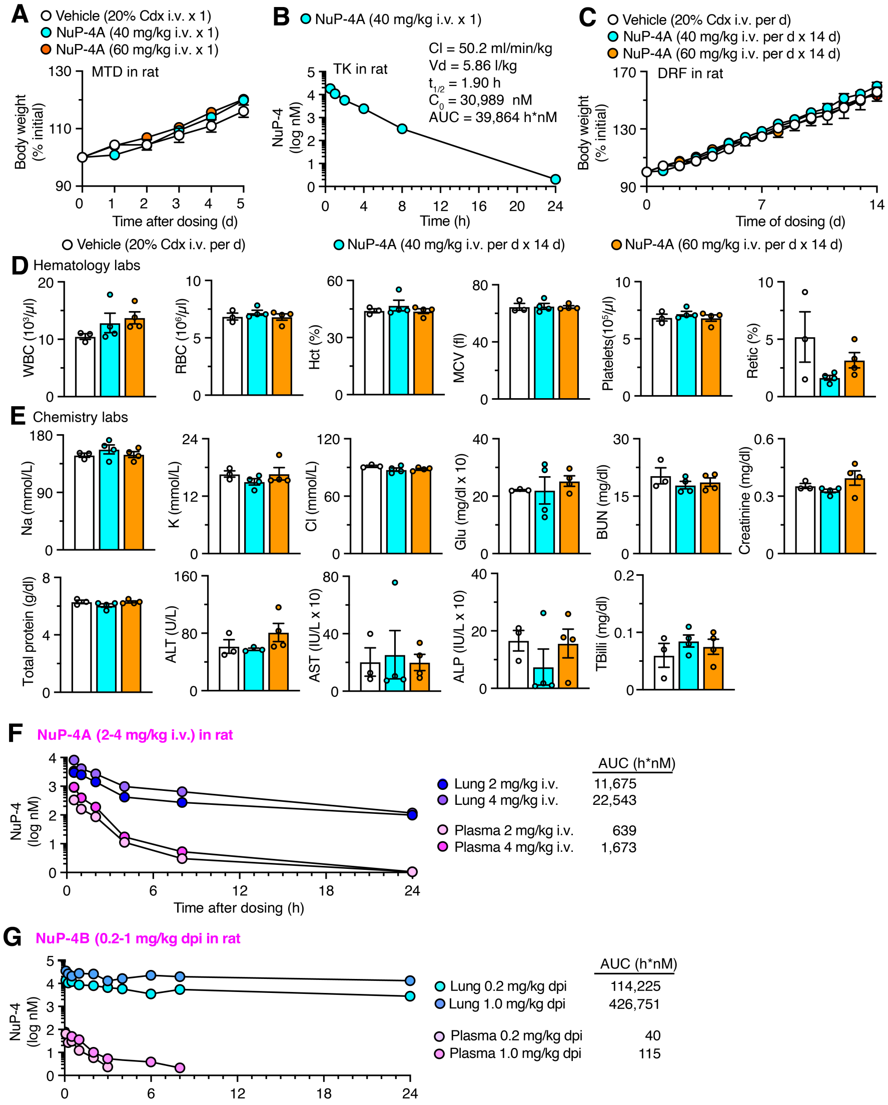


**Supplemental Fig. 1. Toxicology and pharmacokinetic (PK) analysis of NuP-4 demonstrates safety in rat toxicology studies. A,** Body weights from maximum tolerated (feasible) dose study limited to 60 mg/kg by compound solubility in excipient 2-hydroxypropyl-β-cyclodextrin (Cdx). **B**, Toxicokinetic (TK) analysis for NuP-4A (assayed as NuP-4) with single dose at 40 mg/kg i.v. **C**, Body weights for dose-range finding (DRF) study at 40 and 60 mg/kg i.v. each day for 14 d. **D**, Hematology lab values for conditions in (C). **E**, Chemistry lab values for conditions in (C). No significant differences were detected from control vehicle (Cdx) condition using ANOVA and Tukey correction. **F,** Lung and plasma levels for NuP-4A (assayed as NuP-4) given iv at 2 and 4 mg/kg with resulting AUC values. **G,** Lung and plasma levels for NuP-4B given by dpi nose-only at 0.2 and 1 mg/kg with resulting AUC values. Values represent mean ± s.e.m. (n=3-4 rats per condition).

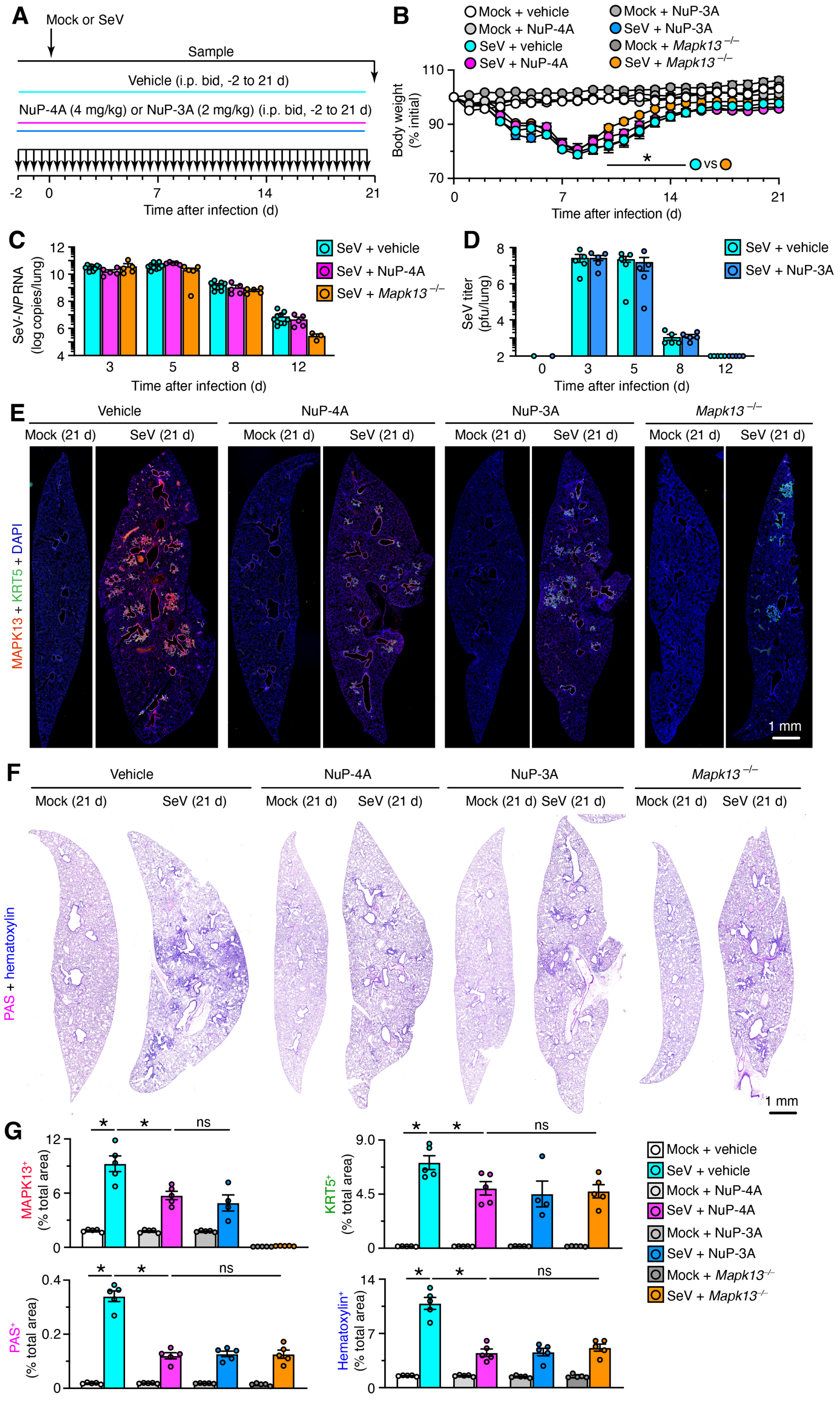


**Supplemental Fig. 2. NuP-4A ip is equivalent to MAPK13-deficiency in preventing lung disease after viral infection. A,** Protocol scheme for NuP-4A versus NuP-3A treatment or *Mapk13*-gene knockout and assessment during acute illness and 21 d after SeV infection or Mock control. **B**, Body weights for conditions in (A). **C,** Levels of viral RNA in lung tissue for conditions in (A). **D**, Corresponding viral titers based on plaque-forming assay for conditions in (C). **E**, Immunostaining for KRT5 and MAPK13 with DAPI counterstaining of lung sections from conditions in (A). **F**, PAS and hematoxylin staining of lung sections for conditions in (A). **G**, Quantitation of staining for conditions in (E,F**)**. Data are representative of three separate experiments with n=5-8 animals per condition in each experiment (mean ± s.e.m.). *$P$ <0.05 using ANOVA and Tukey correction.

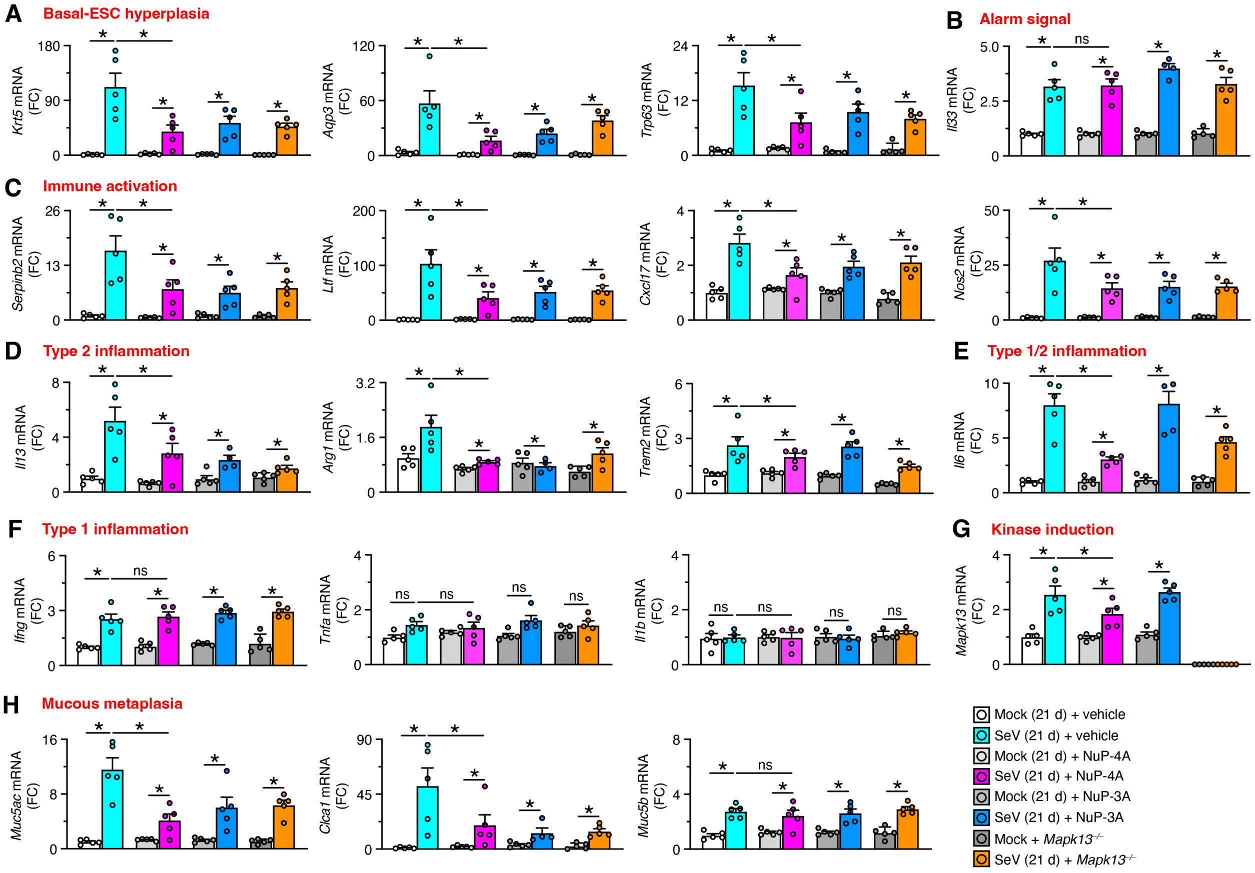


**Supplemental Fig. 3. NuP-4A ip is equivalent to MAPK13-deficiency in preventing muco-obstructive lung disease at 21 d after viral infection based on mRNA biomarkers. A-H,** Lung levels of mRNA biomarkers for indicated disease endpoints for conditions in Supplemental Figure 2A. Data are representative of three separate experiments with n=5 animals per condition in each experiment (mean ± s.e.m.). $*P$ <0.05 using ANOVA and Tukey correction.

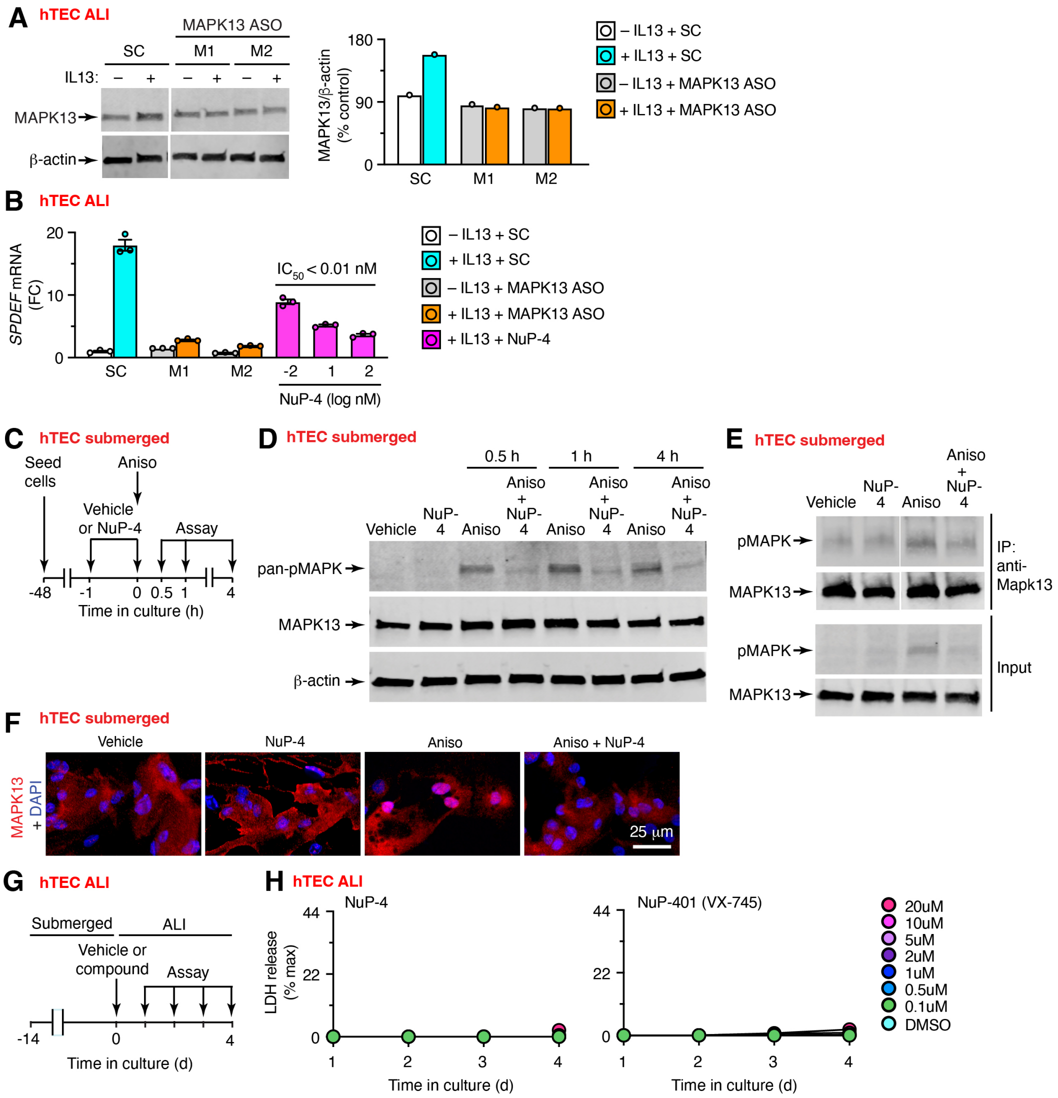


**Supplemental Fig. 4. NuP-4 control over MAPK13 in human basal-ESCs**. **A,** Western blot for MAPK13 levels from MAPK13 ASO treatment conditions in Figure 8D. **B,** Levels of mucous metaplasia (marked by *SPDEF* mRNA expression) for scrambled control (SC), *MAPK13* MAPK13 ASO, or NuP-4 treatment conditions in Fig. 8D,E. **C**, Protocol scheme for hTEC submerged culture designed to preserve basal-ESCs and assess MAPK13 activation in response to MAPK2K2/3 activator anisomycin (Aniso). **D**, Western blot for pan-phospho-MAPK, MAPK13, and control β-actin for Aniso with or without NuP-4 treatment as diagrammed in (C). **E**, Western blot for MAPK13 immunoprecipitation for 1-h conditions in (D). **F**, Representative immunostaining for MAPK13 with DAPI counterstaining for conditions in (E). **G,** Protocol scheme to assess cell toxicity of compounds in hTEC ALI culture conditions. **H,** Levels of cytotoxicity marked by LDH release after treatment with NuP-4, NuP-401, or vehicle for conditions in (G). Data is derived from a single subject (mean ± s.e.m.) that is representative of 3 subjects.

**Supplemental Table 1. Characteristics of tissue donor groups.**

| | Donor (n=11) | Asthma (n=10) | COPD (n=10) |
|---|---|---|---|
| Age | 43 ± 18.4 | 43.8 ± 17.6 | 61.8 ± 5.7 |
| Sex (M/F) | 7/4 | 5/5 | 4/6 |
| Race (W/B/H) | 7/0/0[1] | 4/4/2 | 9/1/0 |
| Pack-year | ND[2] | None[3] | 48.5 ± 27.0 |
| Years quit | ND | ND | 10.3 ± 8.8 |
| FVC % pred | ND | ND | 67.5 ± 11.4 |
| FEV1 % pred | ND | ND | 21.0 ± 4.1 |
| FEV1/FVC ratio | ND | ND | 0.25 ± 0.04 |

[1]Race data missing for some donor subjects.

[2]Two deceased donor subjects had a history of tobacco smoking 10 pack-years and 15 pack-years, but smoking data was not available for all subjects.

[3]Two deceased asthmatics had less than a 5 pack-yr history of tobacco smoking.

Abbreviations: ND, not determined.

**Supplemental Table 2**. Sequences of DNA primers and probes for determining levels of SeV RNA in real-time qPCR assays.

| Target gene | Type | Sequence |
|---|---|---|
| SeV-*NP* | F[1] | 5'-GGCGGTGGTGCAATTGAG-3' |
| | R | 5'-CATGAGCTTCTGTTTCTAGGTCGAT-3' |
| | P | 5'-AGCTCTAGACAATGCC-3' |

[1]Abbreviations: F, forward primer; R, reverse primer; P, MGB probe.

**Supplemental Table 3.** Antibodies for immunostaining mouse and human tissues and cells.

| **Target Protein** | **Antibody Type** | **Vendor** | **Catalogue #** |
|---|---|---|---|
| F4/80 | Rabbit mAb | Cell Signaling | 70076 |
| IL-33 | Goat mAb | R&D Systems | AF3626 |
| KRT5 | Rabbit pAb | Abcam | ab53121 |
| KRT5 | Chicken pAb | Biolegend | 905904 |
| MAPK13 | Rabbit pAb | R&D Systems | AF1519 |
| Phospho-MAPK13 | Mouse mAb | Cell signaling | 9216 |
| MUC5AC | Mouse mAb (45M1), biotinylated | Thermo Scientific | MS-145-B |
| MUC5AC | Mouse mAb (45M1) | Thermo Scientifc | MS-145-P |
| MUC5B | Rabbit pAb | Abcam | ab87276 |
| NOS2 | Rabbit pAb | Abcam | ab3523 |
| SPC | Rabbit pAb | Abcam | ab90716 |
| MBP | Mouse mA BMK-13 | Fisher | NBP1421401M |

[1]Abbreviations: mAb, monoclonal antibody, pAb, polyclonal antibody.

**Supplemental Table 4**. Sequences of DNA primers and probes for real-time qPCR assays in mouse tissue samples.

| Target Gene | Type | ID/Sequence |
|---|---|---|
| *Arg1* | | Mm00475988_m1 (ThermoFisher Scientific) |
| *Aqp3* | | Mm.PT.58.13308206 (Integrated DNA Technologies) |
| *Clca1* | F[1]<br>R<br>P | 5'-ACCGGCTGCCGCTAAAGAGCTTGAG-3'<br>5'-AGACCATTGTTCTGAACCTGATCCGAAG-3'<br>5'-AGCTGTCCAAAATGACAGGAGGCCTGCAGACATA-3' |
| *Cxcl17* | | Mm.PT.58.28640067 (Integrated DNA Technologies) |
| *Gapdh* | | Mm.PT.39a.1 (Integrated DNA Technologies) |
| *IFNg* | | Mm.PT.58.41769240 (Integrated DNA Technologies) |
| *Il1b* | | Mm.PT.58.41616450 (Integrated DNA Technologies) |
| *Il6* | | Mm.PT.58.10005566 (Integrated DNA Technologies) |
| *Il13* | F<br>R<br>P | 5'-GGAGCTGAGCAACATCACACA-3'<br>5'-CACACTCCATACCATGCTGCC-3'<br>5'-CCAGACTCCCCTGTGCA-3' |
| *Il33* | | Mm00505403_m1 (ThermoFisher Scientific) |
| *Krt5* | | Mm.PT.58.41573083 (Integrated DNA Technologies) |
| *Ltf* | | Mm00434787_m1 (ThermoFisher Scientific) |
| *Mapk13* | F<br>R<br>P | 5'-GGAGCTACCCAAGACCTACCT-3'<br>5'-TGTCCGCTTGTCGATGGCCGA-3'<br>5'-GCGCACGTCGGCA-3' |
| *Muc5ac* | F<br>R<br>P | 5'-TACCACTCCCTGCTTCTGCAGCGTGTCA-3'<br>5'-ATAGTAACAGTGGCCATCAAGGTCTGTCT-3'<br>5'-TATACCCCTTGGGATCCATCATCTACA-3' |
| *Muc5b* | F<br>R<br>P | 5'-CTTTCACCCTCAGGAACACGAT-3'<br>5'-TTCGAGGATTATACAGTTCAAAGCA-3'<br>5'-TGAAGGACAAGGTGTGGAGATT-3' |
| *Nos2* | | Mm.PT.58.43705194 (Integrated DNA Technologies) |
| *SerpinB2* | | Mm.PT.58.13584177 (Integrated DNA Technologies) |
| *Tnfa* | | Mm.PT.58.12575861 (Integrated DNA Technologies) |
| *Trem2* | | Mm.PT.58.7992121 (Integrated DNA Technologies) |
| *Trp63* | | Mm.PT.58.11081628 (Integrated DNA Technologies) |

[1]Abbreviations: F, forward primer; R, reverse primer; P, MGB probe.
